# Motion Blur Microscopy

**DOI:** 10.1101/2023.10.08.561435

**Authors:** Utku Goreke, Ayesha Gonzales, Brandon Shipley, Madeleine Tincher, Oshin Sharma, William Wulftange, Yuncheng Man, Ran An, Michael Hinczewski, Umut A. Gurkan

## Abstract

Imaging and characterizing the dynamics of cellular adhesion in blood samples is of fundamental importance in understanding biological function. *In vitro* microscopy methods are widely used for this task, but typically require diluting the blood with a buffer to allow for transmission of light. However whole blood provides crucial mechanical and chemical signaling cues that influence adhesion dynamics, which means that conventional approaches lack the full physiological complexity of living microvasculature. We propose to overcome this challenge by a new *in vitro* imaging method which we call motion blur microscopy (MBM). By decreasing the source light intensity and increasing the integration time during imaging, flowing cells are blurred, allowing us to identify adhered cells. Combined with an automated analysis using machine learning, we can for the first time reliably image cell interactions in microfluidic channels during whole blood flow. MBM provides a low cost, easy to implement alternative to intravital microscopy, the *in vivo* approach for studying how the whole blood environment shapes adhesion dynamics. We demonstrate the method’s reproducibility and accuracy in two example systems where understanding cell interactions, adhesion, and motility is crucial—sickle red blood cells adhering to laminin, and CAR-T cells adhering to E-selectin. We illustrate the wide range of data types that can be extracted from this approach, including distributions of cell size and eccentricity, adhesion durations, trajectories and velocities of adhered cells moving on a functionalized surface, as well as correlations among these different features at the single cell level. In all cases MBM allows for rapid collection and processing of large data sets, ranging from thousands to hundreds of thousands of individual adhesion events. The method is generalizable to study adhesion mechanisms in a variety of diseases, including cancer, blood disorders, thrombosis, inflammatory and autoimmune diseases, as well as providing rich datasets for theoretical modeling of adhesion dynamics.

## Main

Cellular interactions, including cell adhesion, migration, and chemotaxis, are important to investigate the mechanisms of diseases including cancer, thrombosis, inflammatory diseases, anemia, and vasculopathy. *In vitro* cellular imaging techniques for hematology generally require the use of an aqueous buffer ^1–3^, which dilutes the sample and allows transmission of light for imaging.^4^ However, buffer solutions replace the original whole blood medium, potentially affecting the biological mechanisms under investigation. For instance, plasma proteins facilitate the interaction of red blood cells with endothelial cells, and red blood cells induce margination of leukocytes and platelets to the vascular wall.^5, 6^ Comprehensively understanding these phenomena requires a physiologically realistic approach that includes the presence of whole blood. Therefore, intravital methods remain the gold standard for studies of important dynamic processes associated with cellular interactions.^7^ Intravital methods include multi- and single-photon microscopy, confocal microscopy, Brillouin spectroscopy combined with light microscopy, lightsheet microscopy, and endomicroscopy.^8–13^ However, intravital microscopy methods are highly costly, and require intensive effort in both setup and analysis, and therefore have limited applicability for the broader research community.^14, 15^ Total internal reflection fluorescence microscopy is a potential *in vitro* alternative for visualizing cellular interactions under whole blood flow (and examples with buffer flow already exist^16, 17^). But fluorophore labeling in whole blood can be challenging. Finally, laser optical imaging has been used for obtaining the number of platelet interactions that occur with a protein substrate in a microfluidic channel,^18^ but this method is limited to a single numeric output (intensity of light scattered over time), and has additional experimental setup complexity. Here, we describe a practical, accessible, and easily adaptable microscopy method that enables real-time imaging of dynamics of cellular interactions under whole blood flow *in vitro* by completely eliminating the need for blood sample dilution. We call our approach *motion blur microscopy* (MBM). MBM leverages blurring to make the cellular interactions that take place at slower velocity scales discernable (Fig. 1). For simplicity, we showcase MBM on protein functionalized surfaces, but MBM also works on endothelialized surfaces. We show that the numbers of adhesive sickle red blood cells (sRBCs) from individuals with sickle cell disease interacting with the endothelial surface are greater than those of healthy RBCs, and diluting the whole blood samples may diminish these interactions or result in aberrant interactions (Fig. 2). Individual cells with a velocity substantially less than the bulk flow (i.e. immobile, adhered cells, or those that are rolling/migrating while in contact with the surface) can be visualized within the whole blood flow. MBM allows *in vitro* analysis of various static and dynamic properties of cellular interactions, all while mimicking key *in vivo* conditions.

**Figure 1.**
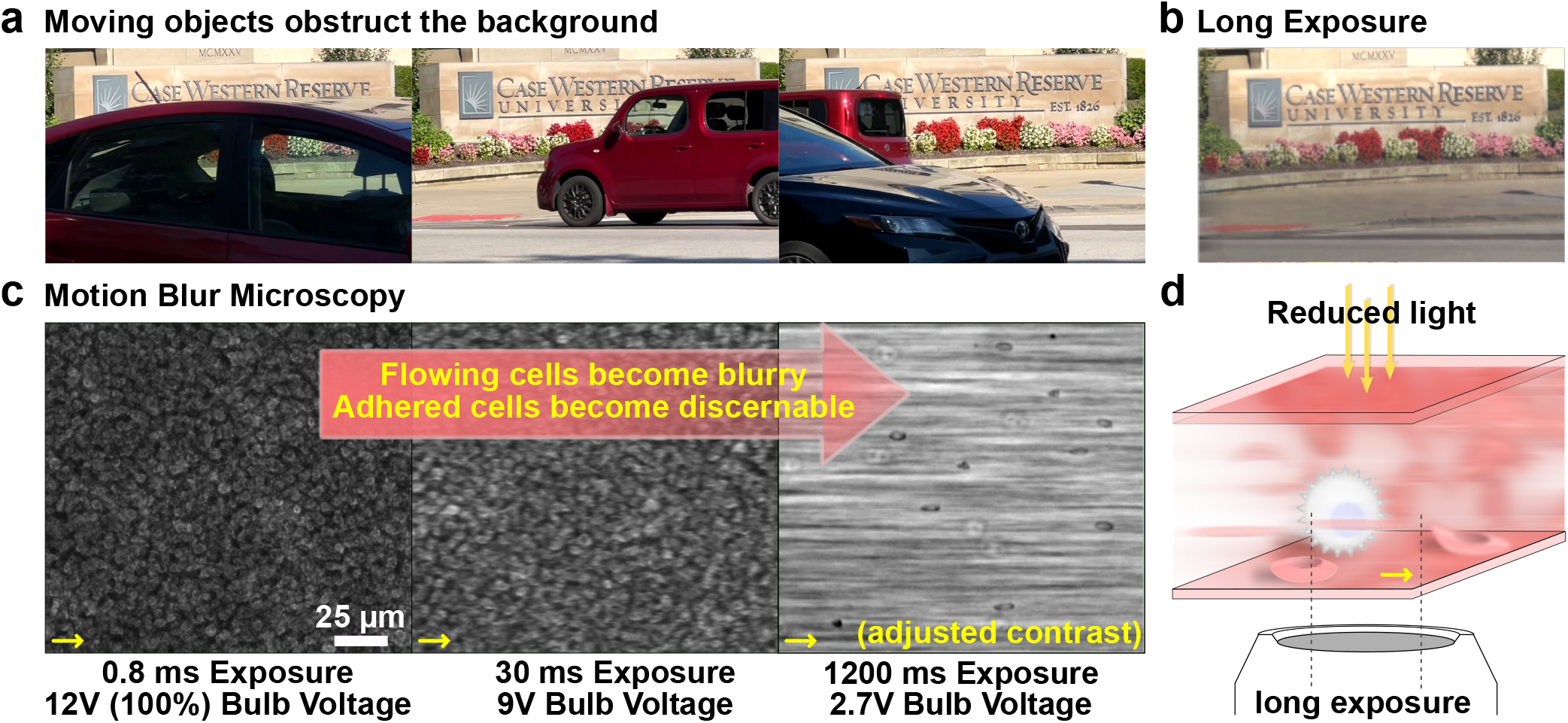
Motion blur microscopy (MBM). **a)** Moving objects in the foreground obstruct the stationary objects in the background. **b)** By adjusting the exposure, moving objects are blurred to obtain a clear view of the background. **c)** The same principle is applied to microscale whole blood flow. Shown are three views of whole blood flow in the same microscopic field with different microscope settings. Increasing the camera exposure (integration time) helps blur the foreground, which consists of non-adhered (flowing) cells. Long exposure results in excess brightness which is then compensated by a reduction of the light source voltage to obtain a clear view of the adhered cells in the background. Yellow arrows show the flow direction. The shear rate of the flow is 50 *s*^*−*1^ which is enough to induce the blur at 1200 ms integration time. **d)** Schematic illustration of the microfluidic channel, with clearly visible adhered cells and blurry flowing cells.

**Figure 2.**
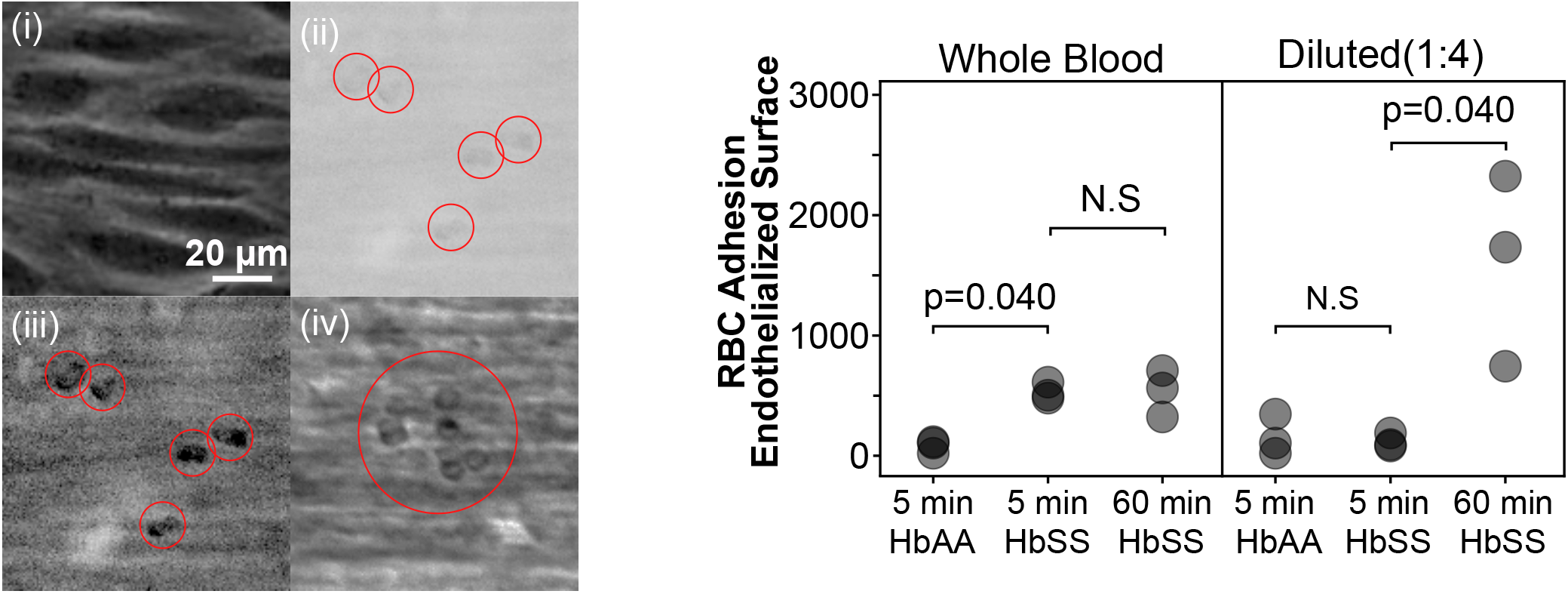
MBM allows capturing cellular interactions on endothelial layers and PBS dilution diminishes the sickle RBC adhesion events. (i) Human umbilical vein endothelial cell (HUVEC) layer on the microfluidic device surface without blood flow. (ii) Original MBM images of sickle red blood cells (RBCs) on the endothelial surface, shown without gray histogram adjustment. (iii) MBM image shown in inset (ii) after stretching the histogram to its limits. (iv) Aberrant cellular aggregations in diluted flow. Under whole blood flow the number of adhered HbSS RBCs was significantly greater than HbAA RBCs. The number of adhered RBCs did not increase after extended flow duration. On the other hand, the dilution of the HbSS whole blood resulted in similar adhesion in HbAA and HbSS samples. Moreover, extended flow duration significantly increased the RBC adhesion through aberrant cellular interactions such as clumping. Red circles denote locations of adhered RBCs. The scale bar applies to all images. P-values are calculated with Mann-Whitney.s

MBM works by reducing the light source and increasing the exposure time, resulting in streaks of flowing cells that generate noisy images. Thus, to identify and analyze adhered cells, an experimenter must spend a considerable amount of time and effort to distinguish cells. While this may not be a difficult task when analyzing individual MBM images, manually analyzing dynamic interactions from MBM videos consisting of hundreds or thousands of frames can be impractical and error-prone. Therefore, we developed an automated machine-learning-based analysis, which can efficiently characterize the dynamics of cellular interactions in MBM videos. The automated analysis is a two-phase system, where phase one identifies groups of pixels in an MBM image corresponding to adhered cells, and phase two classifies these groups by cell type (Fig 3a). The phase one task is completed using a machine learning segmentation network. The phase two task is completed by classifying cells by their size, or by using a machine learning classification network, depending on the complexity of the system under study. Fig 3b shows examples of adhered regions that one might expect the automated analysis to classify. Fig 3b(i) and (ii) correspond to regions of interest, containing sRBCs and CAR-T cells respectively, and the automated analysis should correctly identify these regions as adhered cells.

**Figure 3.**
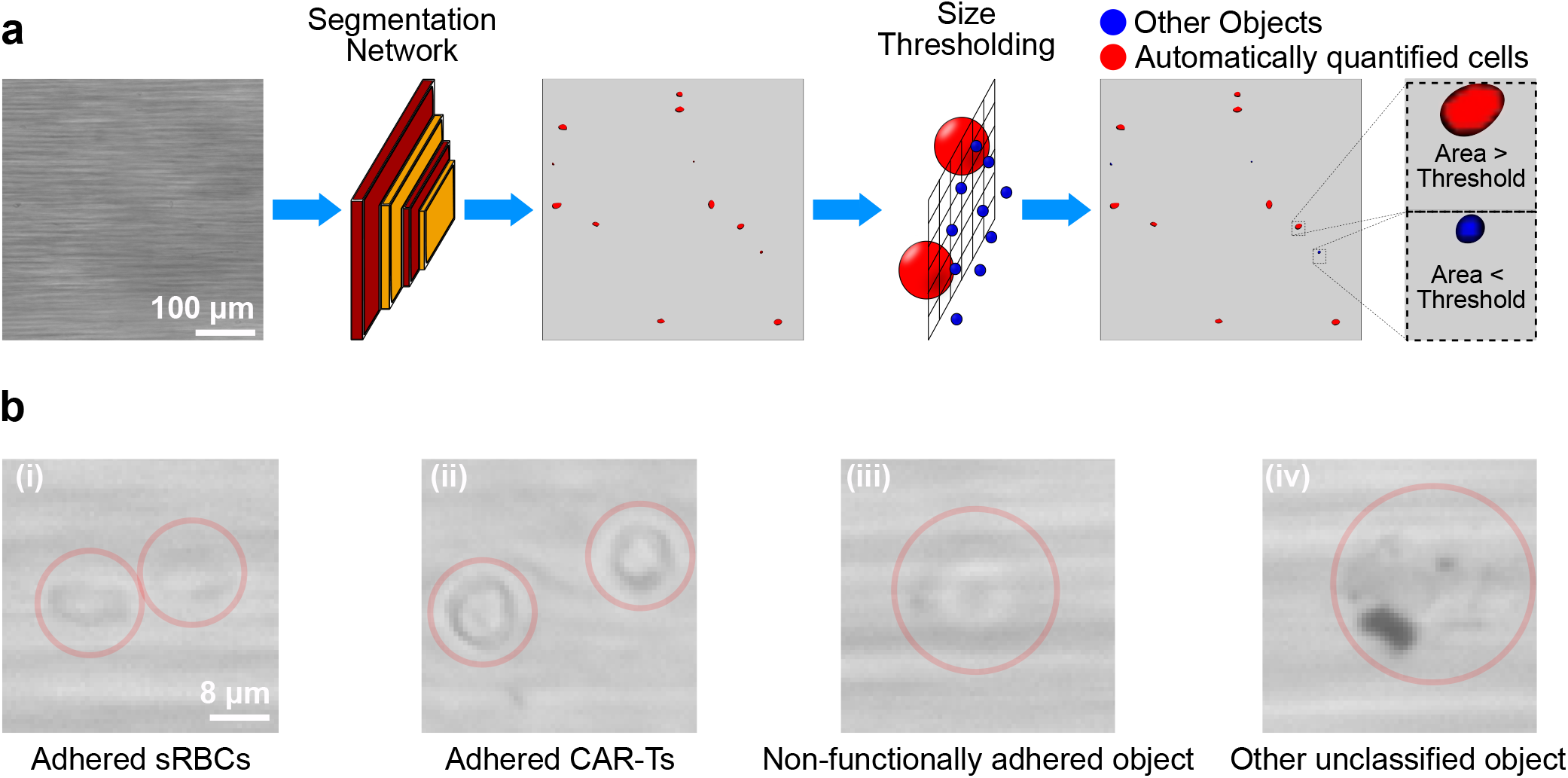
Cartoon diagram of automated analysis pipeline and example objects. (a) The pipeline is conducted in two distinct phases. In the first phase, groups of adhered pixels that may correspond to an adhered cell are identified. In phase two, these groups are classified by cell type. (b) Four examples of adhered objects that might appear in MBM images. (i, ii) show cells of interest, while (iii, iv) show non-relevant adhered regions.

Fig 3b(iii) depicts a typical object adhered to the surface that is not protein functionalized, and Fig 3b(iv) shows other stationary objects that are not cells. The automated analysis pipeline distinguishes these non-functionally adhered objects and other non-cell objects from the adhered cells of interest.

With our automated analysis, we can process MBM images in an accurate and high throughput manner. Adhered cells can be identified, and morphological features, such as the size and eccentricity of each individual cell can be extracted. MBM videos enable studying the dynamics of cells on the individual level, and allow us to determine kinetic properties like adhesion durations, and average velocities (for the case of cells that roll or migrate while on the surface). The individual cell data can be aggregated to produce group statistics, such as distributions of morphological features and dynamic quantities, or mean squared displacements. Importantly, the method allows us to identify and analyze the properties of hundreds of thousands of cells. These data can be used to help us understand cellular adhesion dynamics, or be used in clinical studies. Since our approach relies on a basic experimental microscopy setup, we anticipate that it will be highly accessible to the broader research community.

Characterizing fundamental cellular interactions on the single-cell level can reveal small sub-populations of cells that initiate pathogenesis. Here, we focus on two diseases where the cellular interactions in the vascular space carry central importance. Analyzing these subpopulations typically requires intravital microscopy, and thus, serves as a means to demonstrate the effectiveness of MBM in challenging applications. The first of these diseases is sickle cell disease, where hemoglobin, the fundamental protein underlying red blood cells’ oxygen transport, polymerizes in a deoxygenated environment. When the hemoglobin polymerizes, a cascade of events can occur, leading to a debilitating disease complication known as a vaso-occlusive crisis. Red blood cells containing polymerized sickle hemoglobin experience physical and chemical changes, resulting in abnormally stiff, dense, and adhesive cells^19–21^. The second disease in focus is malignant solid tumors where T-cell migration is crucial for immunotherapy of this type of malignancy. Many chimeric antigen receptor T-cell (CAR-T) therapies are under development for solid tumors, and they hold great promise for treating refractory cancers. Understanding the migration of T cells, specifically CAR-T cells, would lead to better design of future cell therapies when diapedesis/migration is an important aspect of the therapy, as in the case of solid tumors. An accessible, reliable, single-cell level *in vitro* method for analyzing CAR-T cell behavior, like MBM, would accelerate overcoming this bottleneck.

To demonstrate MBM and put its validity to the test, three distinct experimental setups were used. One experimental setup used a laminin functionalized channel to observe sickle red blood cells under flow. A second experimental set-up used an E-selectin functionalized channel to observe CAR-T cells under flow. The final experimental set-up used a P-selectin functionalized channel to observe a combination of red blood cells and CAR-T cells under flow. We include this final set-up as an additional demonstration of the flexibility of MBM, and the details of its analysis are in the Supplementary Information (SI). In this work, we aim to accomplish three main goals. One, we show that MBM with automated analysis is accurate, insofar as it can correctly classify cells for a variety of inputs in a reproducible manner. Two, we show that relevant physical properties of identified cells can be found. Three, we show that the identified cells and their properties constitute a data set with practical uses, ranging from studying cellular adhesion mechanics to aiding in clinical studies.

### Overview of MBM with automated analysis

Here we summarize the complete process for analyzing blood samples using MBM (full details can be found in the Methods and SI). Given a cell type (or types) of interest, we functionalize a microchannel with suitable adhesion proteins. Blood samples sent through the microchannels are then imaged to produce either individual MBM images or videos. Cells of interest adhere to the functionalized protein surfaces, and become visible against the blurred foreground of non-adhered flowing cells. The MBM images are analyzed via a two-stage automated process schematically illustrated in Fig. 3a. The first phase consists of a segmentation neural network, which labels pixels from MBM images corresponding to adhered objects, and groups together neighboring labeled pixels. In phase two of the analysis, groups of labeled pixels are then classified by cell type using either a size threshold (in applications where size is a sufficient criterion) or a specially trained classification neural network (in more complicated scenarios). After all of the cells in an MBM image or MBM video have been identified, we then generate data for each identified cell. Morphological properties like the size and eccentricity of each cell can be extracted, as well as dynamical properties such as adhesion duration or mean velocity. Using the automated analysis, we were able to extract properties of individual cells for 177 sickle red blood cell / laminin images, one CAR-T / P-selectin video with 999 frames, and one CAR-T / E-selectin video with 499 frames, all together containing hundreds of thousands of adhesion events.

### MBM-based cell classification is reproducible and accurate

In cases where only a single cell type is expected to be visible in an MBM image, we demonstrate that we can distinguish groups of adhered pixels identified by the phase one segmentation network from debris and other artifacts using morphological properties of the group.

In Fig 4, we show joint probability densities of the size and eccentricity of all groups of adhered pixels identified by the segmentation network for two collections of MBM images: sickle red blood cell adhesion to laminin (Fig 4a) and CAR-T adhesion to E-selectin (Fig 4b). The histograms on the top and right edges of the figures are marginal distributions of either the size or eccentricity alone. The joint probabilities reveal three distinct classes of objects: small size / high eccentricity, small size / zero eccentricity, and large size / high eccentricity. The last category corresponds to sickle red blood cell or CAR-T cells, and hence we can use a size threshold (indicated by dashed vertical lines) to distinguish cells from other adhered objects (i.e. debris). Each group of pixels above the threshold is classified as a cell.

**Figure 4.**
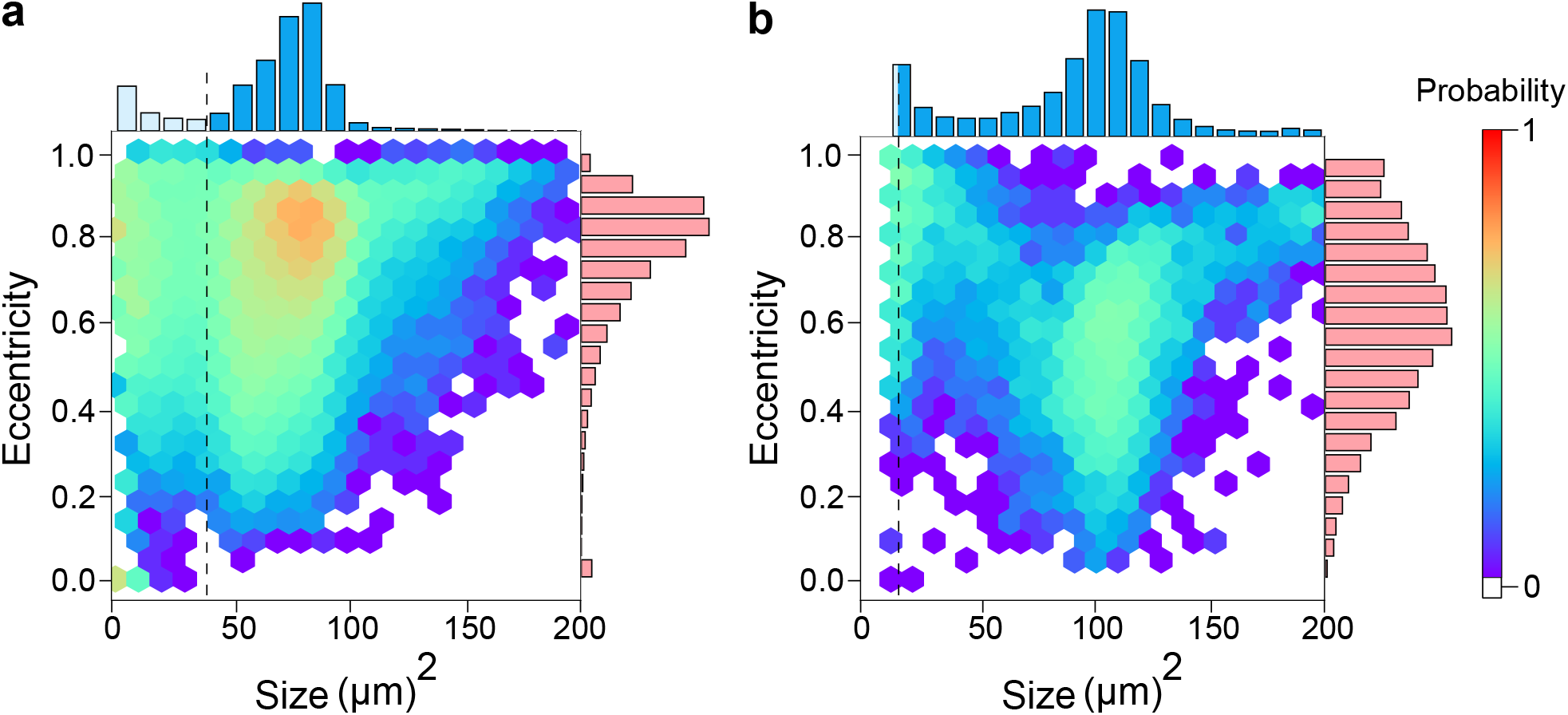
Size and eccentricity distributions under whole blood flow. Joint probability distributions for the size and eccentricity of **a)** objects adhered to laminin functionalized channels from 174 images (*N* = 162, 207) and **b)** objects adhered to E-selectin functionalized channels from 1 video (*N* = 5919). Marginal distributions of size and eccentricity are shown on the top and right axes respectively. Dotted vertical lines indicate the size threshold used for classification, with objects above the threshold corresponding to sickle red blood cells (in panel a) or CAR-T cells (in panel b).

To validate the inter-experimenter reproducibility of this classification scheme, two researchers replicated five consecutive experiments each. The experimenters used a single tube of blood collected from a homozygous sickle-cell disease subject and analyzed the number of sickle cells using the MBM approach five times each over a fifteen minute period per experiment. The results did not show a significant difference between the experimenters (Fig 5a, p= 0.934, two-way ANOVA with replication). Furthermore, the coefficient of variation within the replications of each experimenter was <25% for all but one data point(Fig 5b), an important precision benchmark for bioanalytical method validation^22^. Collectively, these results provide reasonable assurance for the acquisition of meaningful results by showing that our experimental procedure and analysis pipeline are precise and independent of the experimenter.

**Figure 5.**
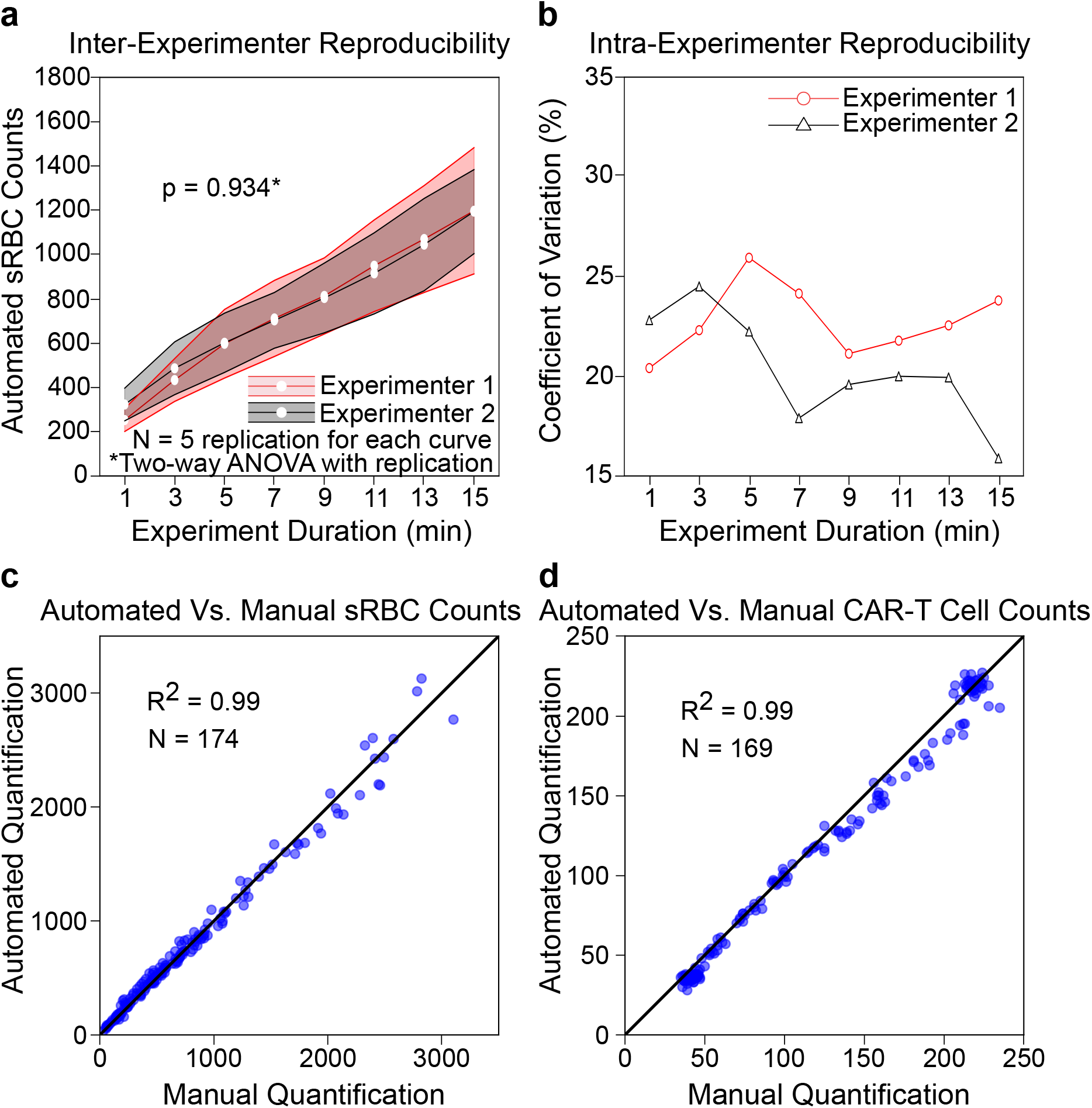
Validation and benchmarking of MBM. **a)** We establish inter-experimenter reproducibility for adhered sickle red blood cell (sRBC) counts for different experimental durations, carried out by two different researchers. Shaded regions around each line are one standard deviation. The cell adhesion results show no significant difference when two different experimenters perform MBM using the aliquots of the same patient sample **b)** We establish the sensitivity of MBM for adhered sickle red blood cell counts. For all but one data point, the coefficient of variation is < 25%, an important benchmark for the precision of bioanalytical methods. Finally, we establish the accuracy of MBM in counting **c)** sickle red blood cells adhered to laminin and **d)** CAR-T cells adhered to E-selectin respectively. There is a strong agreement between automated and human counts in both cases, as indicated by the *R*^2^ value being close to one.

Finally, we tested the accuracy of the procedure in counting cells, by comparing manual human and automated counts of adhered cells in various MBM images for both sickle red blood cell adhesion to laminin (Fig 5c, *N* = 174) and CAR-T adhesion to E-selectin (Fig 5d, *N* = 169). The pipeline performed very well, with an *R*^2^ value of 0.99 for both the sRBC and CAR-T cases. Importantly, the counts remained accurate even as the number of adhered cells becomes large in an individual image.

### MBM can provide high-throughput single-cell dynamic data

The effectiveness of MBM at analyzing adhered cells in static images generalizes to videos, allowing us to characterize the dynamics of adhered cells on protein-functionalized surfaces. To facilitate this, we combined the classification procedure described above (applied to each frame of the video) with a cell tracking algorithm, described in detail in the SI. The tracking analysis distinguishes the motion of adhered cells between sequential frames from new adhesion events, and quantifies the total time spent by a cell on the surface before detachment.

We show eight representative cell trajectories in Fig 6a&b, four for sickle red blood cell adhesion to laminin, and four for CAR-T cell adhesion to E-selectin. We highlight three distinct types of adhesion events: adhesion events with large displacements, adhesion events with small displacements in the same direction as the flow, and adhesion events with small displacements in the opposite direction of the flow. These trajectories are generated for each adhered cell in an MBM video, giving us a high-throughput approach to collect dynamical information about large numbers of cells.

**Figure 6.**
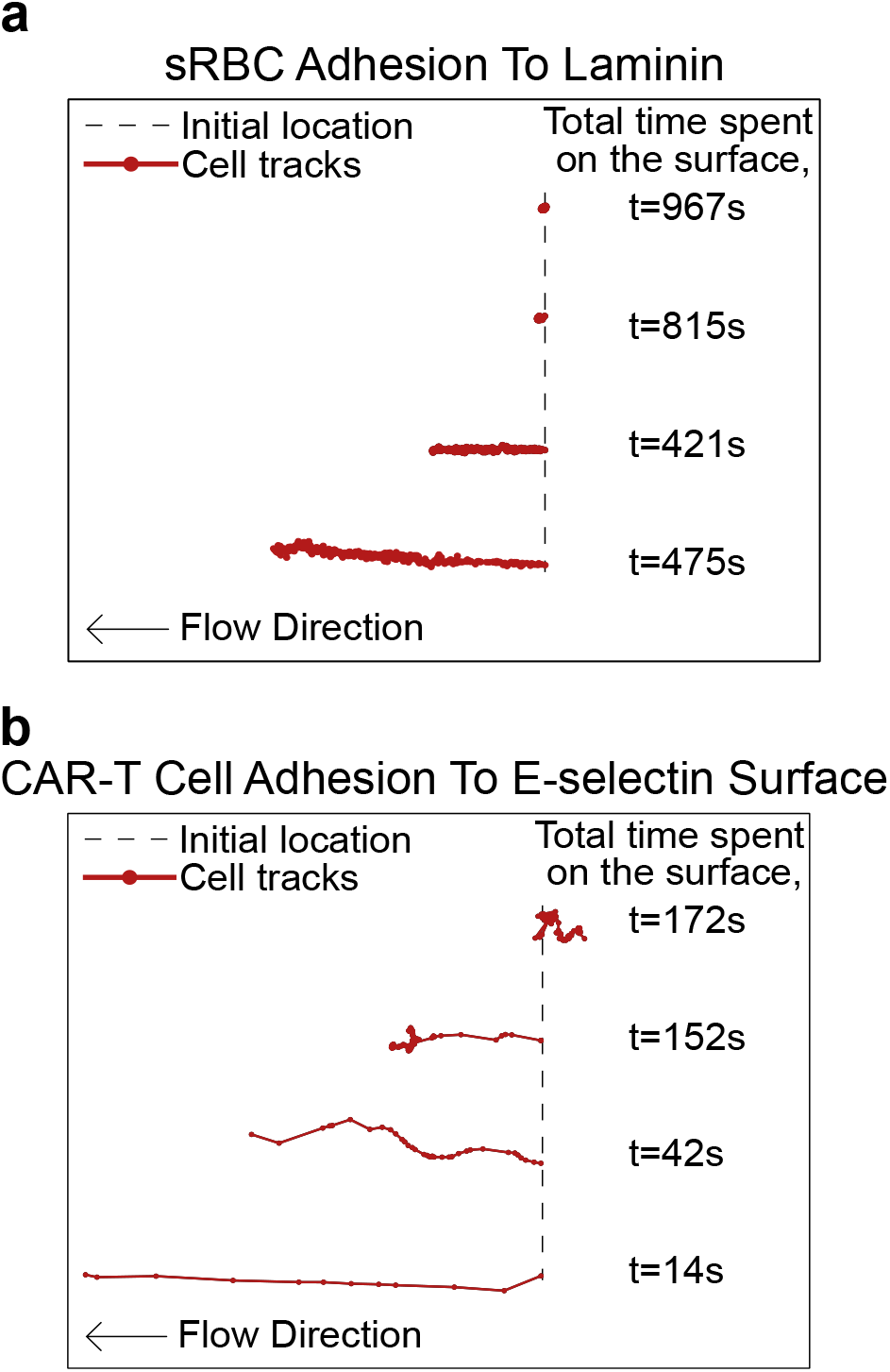
Example adhered cell trajectories. Shown are **a)** sickle red blood cell (sRBC) motion on laminin and **b)** CAR-T cell motion on E-selectin. Generally speaking, the longer a cell is adhered, the smaller the displacement of the cell in the direction of the flow. CAR-T cells tend to have higher motility relative to sickle red blood cells.

The adhesion duration distributions in Fig 7 are one example of the dynamical data that can be compiled through our method, corresponding to thousands of individual trajectories. Fig 7a shows sickle red blood cell adhesion to laminin, and Fig 7b shows CAR-T cell adhesion to E-selectin. In both cases there is a peak in the distribution at low adhesion durations, corresponding to a large number of rapid attachment/detachment events, but also a non-trivial proportion of long-lived trajectories. At intermediate times, both plots are approximately power-law. At long times, the sickle red blood cell case continues the power-law trend, while CAR-T cells experience more rapid decay. The power law trend also applied to HbAA RBCs from healthy donors without known hemoglobinopathies for short-lived RBC adhesion events and was independent of oxygenation conditions for both HbSS and HbAA RBCs (Fig. SI5). The longest adhesion durations (hundreds of seconds) occur infrequently, but the number of events collected by MBM is sufficiently large to resolve these rare cases. Adhesion duration distributions are essential raw data for biophysical modeling of bond dynamics between cells and surface proteins under flow conditions, providing a valuable starting point for future studies.

**Figure 7.**
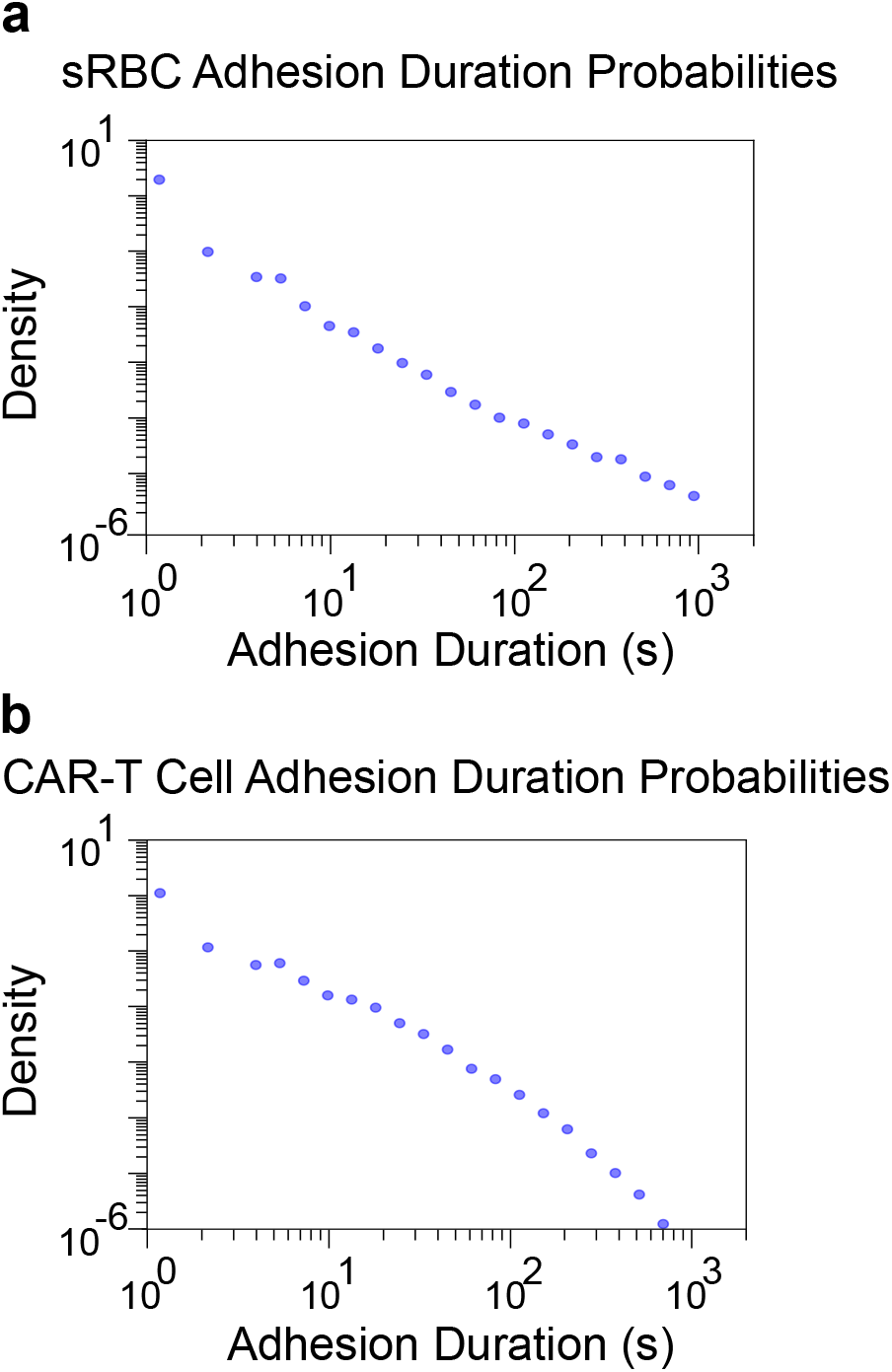
Probability distributions of adhesion durations. Shown are **a)** sickle red blood cell (sRBC) adhesion to laminin (*N* = 14229) and **b)** CAR-T cell adhesion to E-selectin (*N* = 7671).

MBM also has the potential of revealing more subtle dynamical relationships that have not been systematically explored, for example how morphological features of cells are correlated with dynamical behaviors at the single-cell level. For example, a longstanding question about sickle red blood cells is whether elongated, irreversibly sickled cells are more adhesive to endothelium than sickle discocytes^23^. In Fig 8 a, we show MBM data for the relationship between average sickle red blood cell eccentricity (calculated over the entire cell trajectory) and adhesion duration to laminin, revealing a significant negative trend (*p* < 0.03, linear regression): cells with longer adhesion durations are slightly less elongated on average. In Fig 8b, we show that such a trend is not statistically significant between average eccentricity and adhesion duration to E-selectin for CAR-T cells (*p* > 0.07, linear regression).

**Figure 8.**
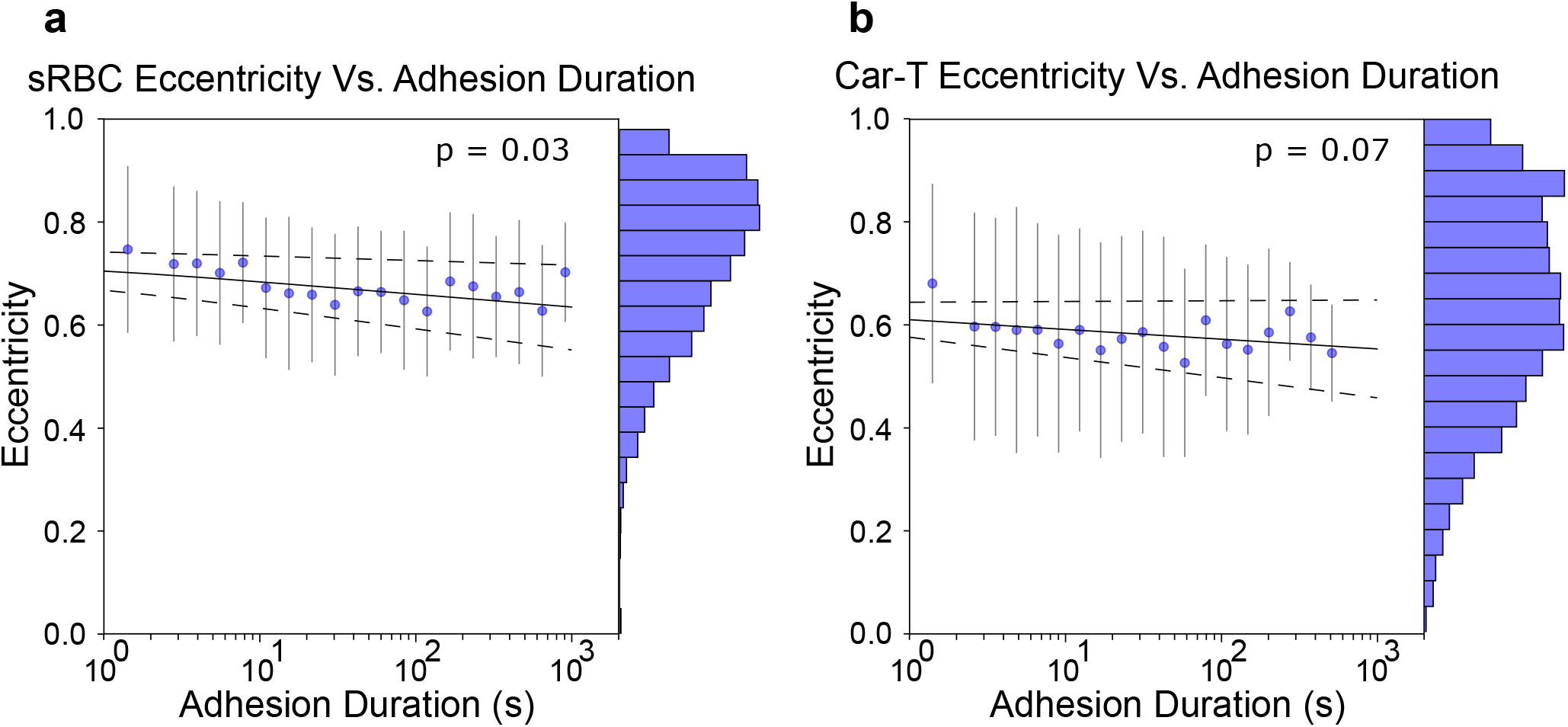
Average eccentricity vs. adhesion duration for (a) sickle red blood cell (sRBC) adhesion to laminin under normoxic conditions (N=14229), and (b) CAR-T cell adhesion to E-Selectin (N=5919). Dotted lines show 95% confidence intervals for the line of best fit. The blue histograms show the distribution of average eccentricities. There is a significant negative relationship between average eccentricity and adhesion duration for sickle RBCs, but not for CAR-T cells (p=0.03, p=0.07, each panel respectively, linear regression). Whiskers for each scatter point are one standard deviation.

We can also investigate how the orientation of the cell motion relative to the flow direction influences the dynamics. For each cell trajectory, the initial and final position (before detachment) defines a net displacement, which divided by adhesion duration gives us an average velocity vector. The left column of Fig 9 shows distributions of the components of this vector parallel to and perpendicular to the flow direction. Moreover, we can correlate the average velocity components with adhesion duration (right column of Fig 9).

**Figure 9.**
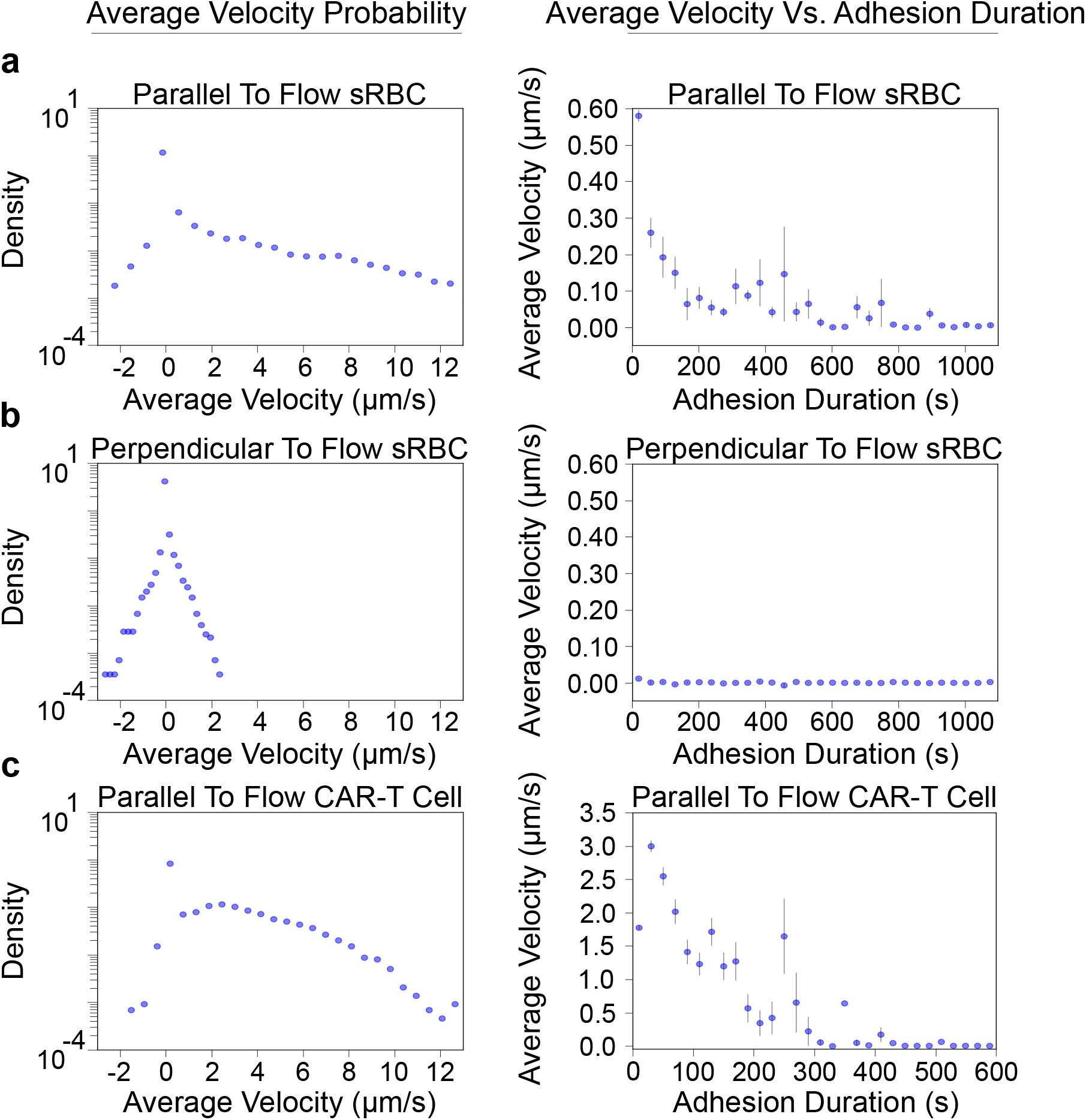
Velocities of cells under whole blood flow. Each row shows a velocity probability distribution (left) and a corresponding average velocity versus adhesion duration plot (right). Three cases are depicted: **a)** Parallel to the flow direction for sickle red blood cell adhesion (sRBC) to laminin (*N* = 14229). The probability distribution shows that a large majority of adhesion events have near-zero velocities. **b)** Perpendicular to the flow direction for sickle red blood cell (sRBC) adhesion to laminin. Because this is the perpendicular to flow direction, we expect the distribution to approach that of a random walk, and the average velocities to be near zero. **c)** Parallel to flow for CAR-T cell adhesion to E-selectin (*N* = 7671). When comparing this row to **a**, we see that CAR-T cells adhering to E-selectin tend to have larger parallel to flow velocities. Perpendicular to flow analysis of CAR-T cells is not shown, but the results are similar to those of **b**.

This analysis reveals a variety of interesting features. The left panel of Fig 9a shows sickle red blood cell velocities on laminin, projected parallel to the flow direction. The distribution is peaked at zero velocity, but has contributions from both positive (with flow) and negative (against flow) velocities. As expected, the distribution is distinctly asymmetric: cells are less likely to move against the flow. The right panel of Fig 9a bins the sickle red blood cells by adhesion duration, and depicts the average parallel velocity for each bin. The shortest adhesion durations have the largest velocities, in agreement with the sample trajectories of Fig 6a. For velocities perpendicular to the flow direction (Fig 9b, left) the asymmetry in the distribution vanishes: average velocities in either the up or down perpendicular direction are equally likely. The large tail at positive velocities that was visible in the parallel distribution also disappears. In order to achieve velocities with magnitudes much greater than 2 *μ*m/s one clearly requires the assistance of flow in the same direction as the cell motion.

Fig 9c analyzes the motility of CAR-T cells adhered to E-selectin, parallel to the flow direction. We noted in Fig 6 that CAR-T cells appeared to be more mobile than sickle red blood cells, and this characteristic is validated in the distribution of average velocities (Fig 9c, left). Relative to the velocity distribution of sickle red blood cells (Fig 9a, left), we see about an order of magnitude higher probabilities at short-intermediate positive velocities. In fact, there is a subsidiary peak in the CAR-T distribution around 2 *μ*s/s, in addition to the zero-velocity peak. This trend is also evident when looking at the correlation of velocity with adhesion duration (Fig. 9c, right). On top of the overall trend of shorter adhesion durations associated with higher velocities, we see that CAR-T cells experience an order of magnitude higher velocity at short-intermediate adhesion durations relative to sRBCs.

Fig. 9 is just one illustration of the versatility of MBM as an *in vitro* platform for blood cell motility analysis, which to date has been exclusively performed with intravital microscopy. Another promising area for exploration is the impact of environmental signals on motility, for example, CD19 activation of CAR-T cells. The majority of CAR-T cells evaluated for B-cell malignancies target CD19. T cells express programmed cell death protein 1 (PD-1) during activation and You et al. used intravital microscopy to show that T cell motility was proportional to PD-1 expression^24, 25^. As an in vitro alternative, we compare CAR-T velocities on E-selectin (parallel to the flow) with and without CD19 activation by using MBM Fig. 10. Fig. 10a shows the velocity distribution for unactivated cells as a control reference, the same distribution as in Fig. 9c but plotted on a linear scale. The velocity bins are colorcoded by different velocity regimes In the absence of CD19 activation, adhered cells fall mainly into relatively immobile populations (the peak around zero velocity, highlighted in purple) and those moving at below 2 *μ*/s (light blue). As expected, the small velocity population consists mainly of cells that adhere at one location, while the mobile cells show a range of trajectory lengths. CD19 activation significantly enhances the mean velocity of CAR-T cells rolling on E-selectin (Fig. 10c, p<0.001, t-test). We also use the same color scale to label and show the trajectories in Fig. 10d&e by their respective velocity regimes. Notably, more complicated motile leukocyte behavior such as crawling can also be captured with MBM (Fig. SI6 and video SI1).

**Figure 10.**
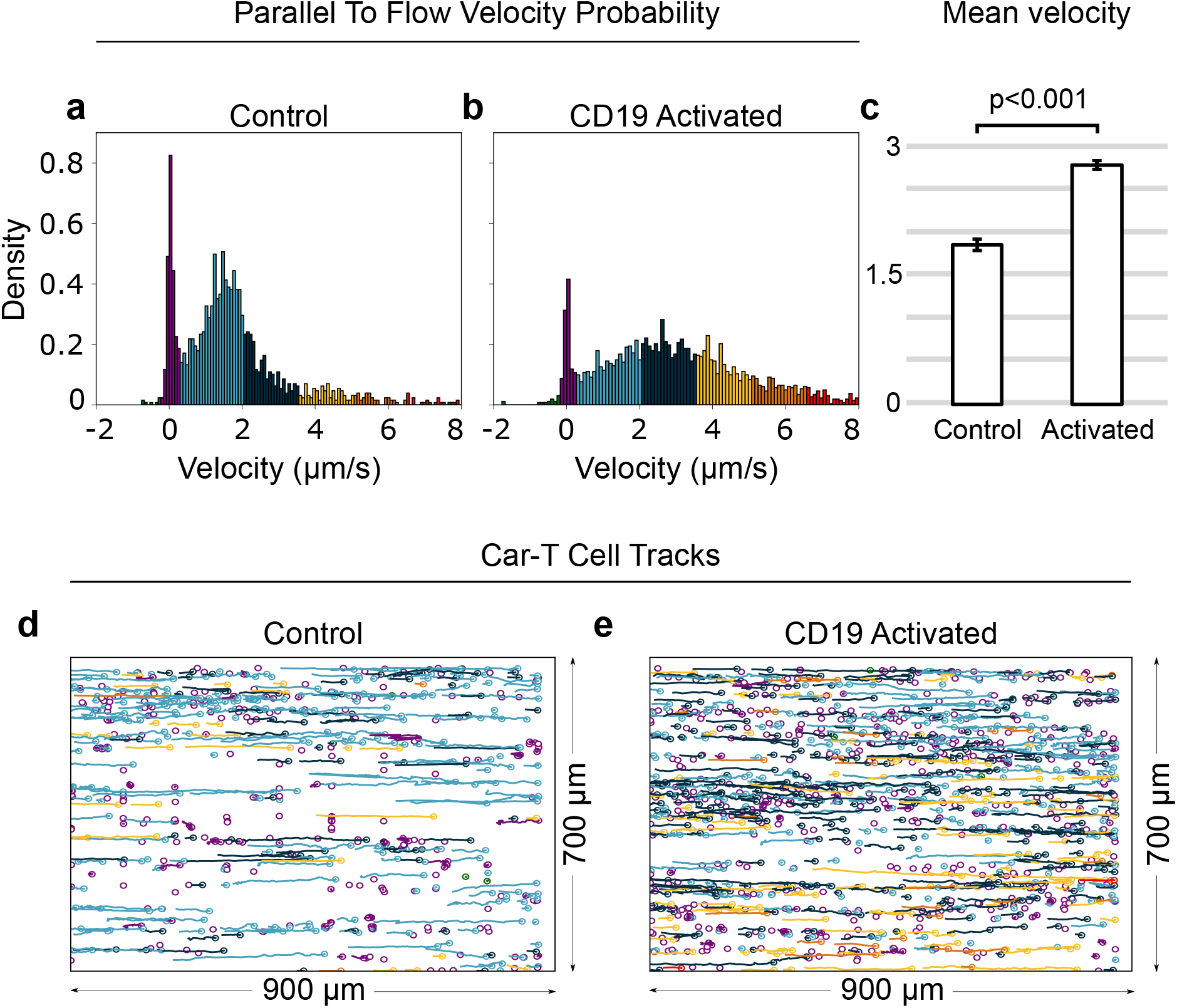
The effect of CD19 activation on the motility of CAR-T cells adhered to E-selectin. a,b) Velocity distribution of CAR-T cells for control (a) and CD19 activated (b) systems, with color coding highlighting different velocity ranges. (c) Activation leads to significantly increased mean rolling velocity among the rolling cells (1.85±0.06 *μ*m/s vs. 2.77±0.06 *μ*m/s, data is mean±95% CI). (d,e) Trajectories of adhered cells for the control (d) and activated (e) cases, labeled by their respective velocity ranges according to the same color scheme as in the distribution panels.

Microfluidics holds great promise for research into the cellular interactions that take place in blood, and as a result various microscopy approaches have been developed which complement microfluidic systems. Yet faithfully recreating *in vivo* conditions in microfluidics is a challenging task, and a bottleneck for its wider adoption. The further a microfluidic study veers from true-to-life conditions, the less physiologically relevant the results become. Therefore, if we are to generate truly meaningful results under *in vitro* conditions, our methods should strive to reproduce the environment of the microvasculature as much as possible. Researchers have developed a variety of approaches to mimic the microvasculature^26, 27^, but the blood—the main constituent with its full complexity—has always been missing in these microfluidic systems. MBM enables researchers to investigate the dynamics of cellular interactions in the presence of blood flow. This means that many of the relevant physical forces and chemical signals which modulate cellular interactions will be present during an experiment. Environmental completeness would be especially crucial when the functional pathways of cells are actively regulated by their environment. One example of such regulation is the influence of the complement and renin-angiotensin systems on leukocyte function^28^. Therefore, we anticipate that MBM will have a formative effect on microfluidic studies. Even so, a new method is only as good as the available tools for analysis. Our machine learning pipeline for MBM can automatically analyze hundreds of thousands of observed adhesion events. Typically, the task of completing this analysis manually would be impossible, as the amount of time required, or the number of people required, would be far too large. Our automated analysis approach is flexible and mitigates errors due to individual human biases in cell counting and classification.

The applications of MBM are wide-ranging for both fundamental investigations into biophysical mechanisms and clinical studies. For example, adhesion duration distributions are crucial for studies of force-dependent binding/unbinding of protein complexes and cell-cell interactions^29^. MBM can complement existing approaches in this area like numerical simulations or force spectroscopy experiments (i.e. atomic force microscopy)^30–36^. MBM could provide a valuable comparison point for bond lifetime simulations, adding extra physiological realism due to the blood flow. Because of its ease of implementation, it is also accessible to a broader research community than atomic force microscopy techniques. On the clinical front, there are a variety of promising applications for specific diseases. In the case of cancer, MBM could help visualize crucial interactions of circulating tumor and endothelial cells with improved physiological accuracy^37^. It can also be utilized to distinguish and isolate rare circulating tumor cells from billions of other cells flowing through a microfluidic channel. For sickle cell disease, MBM could serve as a means to understand how sickle cells behave under changes in blood pressure, flow rates, and viscosity, as well as concentrations of relevant constituents in whole blood. Furthermore, the concentration of adhered sickle red blood cells, as well as the morphological characteristics of the cells may be used to monitor if a sickle cell disease patient is undergoing a flare-up, or if they are in an asymptomatic diseased state. Other possible disease contexts where MBM may be useful are solid tumor malignancies. MBM may help us characterize the motility of immune cells including T-cells and macrophages with great physiological relevance. As a final example, MBM could be used to further our understanding of autoimmune diseases, such as rheumatoid arthritis, where leukocyte recruitment is an important part of the pathology. For example, MBM can be used in tandem with “joint on a chip” systems, where complex interactions among blood cells, endothelium and skeletal components are studied with *in vitro* experiments^38^.

Aside from clinical studies, MBM with automated analysis has the ability to efficiently gather large quantities of data that may provide an ideal basis for theoretical modeling of cellular adhesion mechanics. For example, the velocity distributions parallel to the flow direction from Fig. 9 qualitatively resemble those of molecular motors that are capable of forward and backward stepping along a cytoskeletal track, which can be described via coarse-grained kinetic models of the underlying biochemical cycle^39^. Similar mathematical approaches could be brought to bear on the cell velocity data, giving us a more complete picture of the cell interactions with the endothelium that give rise to these kinds of dynamics. The excellent statistics of the data set is crucial in this regard, since in principle it enables us to distinguish between competing models.

As with any experimental approach, there are also limitations. In sufficiently complex scenarios where large densities and/or multiple cell types adhere to the surface, extracting single-cell dynamical information may become more difficult. Large densities, with many overlapping adhered cells, could impede the video analysis in determining if two cells in consecutive frames are the same or not. Overlaps were rare in the examples we investigated, allowing for straightforward extraction of cell trajectories. In physical systems where this would be an issue, however, more sophisticated methods could be incorporated into MBM analysis workflow for tracking cells^40, 41^. Similarly, both the sickle red blood cell and CAR-T datasets we focused on in the main text involve a single adhered cell type, which could be identified reliably using a size threshold. In cases with multiple cell types of similar sizes, we would have to employ a more involved classification procedure, using other morphological characteristics of the cells. The SI shows results from one such example, where a convolutional neural network was trained to reliably distinguish CAR-T cell adhesion to P-selectin from red blood cells, which also adhere to the surface. Thus we believe all these limitations may be substantially addressed with additions to the data analysis approach. Extended exposure time inherently offsets the eccentricity estimations of the interacting cells. The magnitude of the offset would depend on the velocity of the cells. Eccentricity estimation of high motility cells including T-cells may be significantly affected by this issue. In a typical scenario, a perfectly round CAR-T cell with a size of 8 *μ*m, moving with a mean velocity of 2 *μ*m/s would travel 2.4 *μ*m during the 1200 ms exposure time of the camera. In this case, the resulting eccentricity estimate would be offset by 0.32. This effect can be accounted for using a velocity-based calibration curve for the major axis *a*, during the eccentricity *e* calculation:

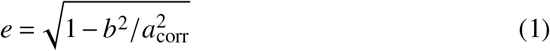

where *b* is the minor axis length and the corrected major axis length *a*_corr_ would be given by:

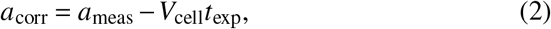

where *V*_cell_ is the velocity parallel to flow, *t*_exp_ is the exposure time of the camera, and *a*_meas_ is the measured major axis length. Strictly speaking, the major axis might not align entirely with the velocity vector, in which case, the calculation would become more complicated. Even still, we believe this simple adjustment should mostly account for changes in eccentricity due to flow velocity. We performed eccentricity offset calibration based on velocity for CAR-T cell adhesion to E-selectin in Figures 4 and 8.

In summary, MBM is a robust, easy-to-implement, high-throughput method to study cell adhesion dynamics in the presence of blood flow. Its flexibility allows for broad deployment in both clinical contexts and studies of fundamental biophysical mechanisms underlying cell adhesion. Combined with automated analysis for cell classification and tracking, it promises to be a general platform for elucidating how interactions with the complex whole blood environment influence and regulate cellular adhesion.

## Methods

### Blood sample collection and microfluidic adhesion assays

Surplus EDTA-anticoagulated whole blood samples were collected under an Institutional Review Board approved protocol (registered at www.clinicaltrials.gov as NCT02824471, “Sickle Cell Disease Biofluid Chip Technology”). The microfluidic platforms were fabricated and functionalized as described previously^42^. For studying the adhesion of HbAA containing red blood cells, unprocessed whole blood samples from healthy donors with no known hemoglobinopathies were used. For studying the adhesion of sickle red blood cells, unprocessed whole blood samples from homozygous sickle cell disease subjects were used and microchannels were coated with laminin (murine, laminin-1). For demonstrating Jurkat or CAR-T motility, the microchannels were coated with either Por E-selectin (human, CD62P and CD62E). For microfluidic surface endothelialization the channels were incubated overnight at 4°C with 10% human fibronectin (Sigma-Aldrich). HUVECs (Lonza) were then seeded into each channel at a concentration of about 10 million cells/*μ*L and channels were cultured under flow conditions created by a peristaltic pump for 2-3 days until confluent. Whole blood samples from subjects with no hemoglobinopathies were first leukodepleted and then premixed with Jurkat or CAR-T cell populations at 40% hematocrit in Hank’s buffer containing calcium to retain the leukocyte activity by replenishing EDTA-depleted Ca++ in blood samples (Fig. SI7). HbSS whole blood was diluted 1:4 with PBS for the dilution experiment. Hypoxic conditions (SpO2 of approximately 83% in the blood sample), using a micro-tube gas exchanger as we described previously^23^. To prevent non-specific cellular interactions, microchannels were incubated with bovine serum albumin at 4°C overnight, and additionally, 250 *μ*l of bovine serum albumin was injected into the microchannels at room temperature with a 5 *μ*l/min flow rate 2 hours prior to the experiments.

### CAR T-cell culture

CAR T-cells were obtained from the Hematopoietic Biorepository and Cellular Therapy core at Case Western Reserve University. These cells manufactured according to ethics and guidelines from University Hospitals Cleveland Medical Center (UHCMC IRB 03-18-01C). Briefly, CD3+ T cells were collected from whole blood samples using the Miltenyi CD3 T cell isolation kit (North Rhine-Westphalia, Germany) and transduced with a CD19-directed CAR vector. Cells were cultured in TexMACS media with IL-7 and IL-15 (Miltenyi Biotec; North Rhine-Westphalia, Germany). Cells were then cryopreserved until experimentation. Upon thawing patient samples, cells were cultured in RPMI 1640 medium with 10% fetal bovine serum, 100 U/mL pen/strep (Thermo Fisher, Waltham, MA, USA), 2 mM glutamax (Thermo Fisher, Waltham, MA, USA). For CD19-activation, CART cells were added to a solution of IL-2 culture media containing CD19+ RAJI cells obtained from American Type Culture Collection (Manassas, VA, USA).

### Motion blur microscopy

Microchannels were visualized with Olympus CellSens software using an Olympus IX83 inverted microscope and QImaging EXi Blue CCD camera with 10x objective (numerical aperture 0.3, pixel area 6.5 *μ*m^2^). To induce motion blur, camera exposure was set to 1.2 seconds. Images and videos of the microchannel surface were saved uncompressed to reduce noise. Frame rates of the videos were kept at (1/1.2) s^*−*1^, which is the maximum frame rate for a 1.2 s exposure time. High integration time was compensated for by adjusting the voltage of the light source to 2.7V (maximum 12V). It should be noted that it is key to adjust the focus map for the surface of interest before the microchannels are injected with the sample. Unprocessed or leukodepleted whole blood was withdrawn into a syringe which then was loaded into a constant displacement syringe pump (NE-1000, New Era Pump Systems Inc.). MBM with automated analysis requires a minimum flow velocity of approximately 150 *μ*m/s in the background to create the minimum particle streak for MBM. Higher background flow velocity yields better distinction of cellular interactions. 50/50 light distribution mode between the camera and the eyepieces was selected, which allows doubling the exposure time at the same brightness level. We performed all the experiments in a dark room. This is important because MBM is a low-light technique and room lights, or sunlight, may introduce non-uniformity to image lightness.. The flow velocity was kept at approximately 500 *μ*m/s for 20 minutes for demonstrating sickle red blood cell adhesion to laminin. For the demonstration of CAR-T cell adhesion to E-selectin, the flow velocity was swept linearly from 500 *μ*m/s to 3500 *μ*m/s in 10 minutes.

### Automated image analysis

To analyze MBM images, we developed an automated analysis procedure with two distinct phases. Phase 1 performs segmentation on MBM images at the pixel level and phase 2 classifies groups of pixels by cell type. After classification, we can perform further analysis on the classified cells (i.e. morphology characterization and trajectory tracking). While we focus on three experimental examples in this paper, our approach can easily be generalized to other systems.

#### Phase 1: pixel segmentation

Phase 1 uses a machine learning network to segment every pixel in an MBM image into one of two categories: adhered or background. Adhered pixels correspond to a pixel of the image belonging to an adhered object, with the remaining pixels falling into the background category.

The segmentation network is based on a modified U-Net architecture from our previous work^43^ (further details are provided in the SI). To train the network, one or more MBM images is chosen as training/validation images. A labeled mask of the training/validation images is created, by manually labelling each pixel as adhered or background. Each labeled training/validation mask, as well as each training/validation image, is then be split into tiles of size 150×150 pixels, which are subsequently resized to 128×128 pixels, as that is the required input of the segmentation network. Once all of the training/validation tiles are compiled, we can then train the segmentation network for our particular task.

For the purposes of this paper, we have trained and used two distinct segmentation networks, one for analysis of sickle red blood cell adhesion to laminin, and one for analysis of CAR-T cell adhesion to E-selectin. The first network was trained on tiles from three MBM images from different sources. The training tiles are drawn from one of the three sources, while the validating/testing tiles are drawn from the other two sources. In total, there were 2163 tiles used for training/validating/testing. The second network was trained on tiles from the frames of three MBM video sources. For each video source, ten evenly spaced frames were extracted from the final 100 frames. The training tiles are drawn from one of the video source frames, while the validating/testing tiles are drawn from the other two video source frames. In total, there were 767 tiles used for training/validating/testing. In both cases a stratified split of 70%/20%/10% training/validating/testing was used.

#### Phase 2: cell classification

The output of the segmentation network is a fully segmented image, where each pixel belongs to one of two classes. In the second phase of the analysis, groups of adhered pixels are extracted and classified. Generally speaking, the cell identification task falls into one of two cases. In the first case, only one adhered cell type is expected to be distinguishable in the MBM image. In the second case, two or more cell types are expected to be distinguishable in the MBM image. For brevity, we will only highlight the single cell type case in the main text, and interested readers can refer to the SI for information on the multiple cell type case.

When only a single cell type of interest is expected to adhere to the functionalized channel, we can classify groups of adhered pixels using a physical property of the group. For example, when we look at the attachment of sickle red blood cells to laminin or CAR-T cells to E-selectin, the vast majority of groups of adhered pixels correspond to one cell type. In this case, we can use a size threshold to distinguish adhered cells from groups of adhered pixels falsely classified by the segmentation network as adhered.

When training both segmentation networks, the size threshold for the classification of groups of adhered pixels was optimized with the objective to produce the closest agreement between human and machine counts of relevant cells. To find the optimum threshold, an array of threshold values can be used to count cells for an array of inputs. The automated cell counts can then be compared to manual cell counts, and the threshold which produces the greatest agreement can be chosen. For single-cell type analysis, two optimum thresholds were found, one for sickle red blood cell adhesion to laminin and one for CAR-T cell adhesion to E-selectin.

In the laminin case, 174 MBM images were analyzed using a trained segmentation network and size thresholds in the range 0-200 px (corresponding to the area of the adhered pixel group), in increments of ten, were used to produce sickle red blood cell counts. Agreement between human and machine counts was checked for each threshold value, and the optimum threshold was found to be 90 pixels (38 *μ*m^2^), where groups of adhered pixels with a size greater than 90 pixels were classified as sickle red blood cells, and groups of adhered pixels with size less than 90 pixels were classified as other. For the CAR-T case, 172 frames from an MBM video were analyzed using a trained segmentation network and size thresholds in the range 0-200 px, in increments of ten, were used to produce CAR-T cell counts. The optimum threshold was found to be 30 pixels (13 *μ*m^2^).

### Post-classification analysis

After phase 1 and phase 2 are completed, the groups of adhered pixels that are positively classified as the cell type(s) of interest can be further analyzed. Properties like the size, eccentricity and position of each individual cell are easily obtainable. These static data can be analyzed individually or aggregated together to produce group statistics. When MBM video is available, we can analyze the dynamics of cells under flow. We define a parameter for each adhesion event called the survival time—the time from when a cell attaches to the surface until it detaches or leaves the frame. With the survival time in hand, as well as the static data for each frame, we can generate dynamic data for each cell. Individual cell dynamics like average velocity, average size, and average eccentricity can be found, or, we can aggregate cells and produce group statistics like mean-squared displacement, or distributions of dynamic quantities.

### Identification rules for tracking moving cells in MBM videos

MBM video analysis brings forth additional challenges beyond those of individual images. In particular, we need to track cells over time, requiring us to determine if cells in consecutive frames are the same cell or not. Two distinct issues arose when making this determination: duplicate counts, and tracking motile cells.

Duplicate counts occur when the segmentation network erroneously segments a single group of adhered pixels as two distinct groups of adhered pixels, whose edges are close, but not touching. To minimize the duplicate counts, we checked to see if the centroids of any two groups of adhered pixels were within 50 pixels (1 px = 0.322 *μ*m) of one another. Also, the shortest distance between the edges of all such pairs was computed, and if this distance was 5 pixels or fewer for any pair, the two groups of adhered pixels were considered to be one. In this case, the two groups of adhered pixels were replaced by the smallest polygon that contains them, known as the *convex hull*. One can imagine this shape as the result of stretching a rubber band over the two groups of adhered pixels and letting it rest taut.

Another case of duplicate counting occurs if phase 2 misclassified an adhered cell as “other” for any amount of frames between two positive classifications. This error would cause the analysis to detect a “new” group of adhered pixels, despite it being the same cell. We avoid this by giving each adhesion event a disappearance allowance. Each cell for an adhesion event was allowed to “disappear” for a number of frames before considering the adhesion event completed. If a cell “disappeared” for a number of frames below the allowance, and then “reappeared”, the adhesion event would continue as if the misclassification never occurred. For sickle red blood cells, we allowed cells to go undetected for up to two frames and still count as the same cell. For CAR-T cells, they could go undetected for up to nine frames. These frame thresholds helped us to not preemptively end adhesion events.

Interacting motile cells are constantly adhering, detaching, and persisting in their adhesion to the surface, and as such, the results of analyzing many frames present a complex picture of many adhesion events happening at various times. Thus, we devised rules by which to determine if a cell in consecutive frames was a new adhesion event or a continuation of a prior adhesion event. If a positively identified group of adhered pixels was within 5 pixels in any direction of a positively identified group of adhered pixels in the previous frame, the group of adhered pixels was classified as a continuation of the adhesion event. Otherwise, the positively identified group of adhered pixels was classified as the beginning of a new adhesion event. Also, if a positively identified group of adhered pixels was within 5 to 30 pixels in the direction of the flow, this group of adhered pixels was classified as a motile adhesion event. These movement rules plus the frame thresholds protected us from splitting a singular adhesion event into two due to any misclassifications by the automated analysis.

## Statistical methods

Statistical analyses were performed using the Microsoft Excel Analysis ToolPak add-in. *p* < 0.05 was chosen to indicate a significant difference.

## Code and data availability

All the code associated with the MBM analysis, as well as a link to the Open Science Framework repository containing the datasets, can be found at: https://github.com/hincz-lab/motion-blur-microscopy.

## Acknowledgments

We express deep gratitude to Paolo F. Caimi at the Case Comprehensive Cancer Center and Cleveland Clinic, Jane Reese Koc at the Hematopoietic Biorepository and Cellular Therapy Core at Case Western Reserve University for sharing the CAR-T cells, and to Paul Tesar at the Case Western Reserve University School of Medicine for sharing the cell culture resources. This work was supported by the National Institutes of Health awards U01AI176469, R42HL162214, R42HL160384, K25HL159358, and National Science Foundation awards 1552782, 1651560, 2112202, 2332121. The authors acknowledge with gratitude the contributions of research participants and clinicians at Seidman Cancer Center (University Hospitals Cleveland Medical Center).

## Authorship Contributions

U.G. and U.A.G developed the idea. U.G., B.S., A.G., M.H. and U.A.G. designed the study. U.G., M.T., O.S., W.W., and Y.M. performed the experiments. U.G., B.S., A.G. and M.H. developed the automated algorithm. U.G., B.S., A.G., M.T., O.S., Y.M, W.W, R.A, M.H. and U.A.G. analyzed the data. U.G., B.S., A.G., M.T., O.S., W.W, M.H. and U.A.G. discussed and interpreted the data. U.G., B.S., A.G., wrote the manuscript, U.G., B.S., A.G., prepared the figures. M.H. and U.A.G. edited the manuscript.

## Supplementary Information

### Motion Blur Microscopy

#### Phase 1 training details

Ideally, the chosen training/validation images are dense in the objects of interest, which will combat any class imbalance during training. Labeled masks for the training/validation images were created by coloring each pixel of the images one of two colors: one color for adhered pixels, and one color for background pixels. This process was completed by first coloring all adhered pixels one color using the software GIMP. Any non-colored pixels were then filled in automatically with a Python script. When resizing tiles from 150×150 pixels to 128×128 pixels, the cubic interpolation method of the Python library OpenCV was used.

To train the segmentation networks (with architecture summarized in Fig. SI1), a custom data generator was created, which loaded tiles into the network in batches of 32 and applied data augmentation. Data augmentations included are rotations of 90, 180, 270, and 360 degrees, horizontal flips, and vertical flips. In total, the augmentations can produce 8 unique orientations of each tile. A stop function was included in the training, which automatically stopped the training after the validation loss had not decreased for a number of consecutive epochs. The weights of the network after the stop function was executed were used as the trained network weights. The loss function used for optimization was a linear combination of the categorical cross-entropy loss (Cat) from the Keras library, and the Jaccard loss (Jac), of the form:

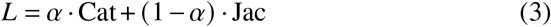

where *α* is a real number in the range [0,1]. The training was completed in both instances using an Adam optimizer. The number of epochs before stopping, *α* value, and learning rate used for training of sRBC adhesion to laminin and CAR-T cell adhesion to E-selectin were [7,0.58,0.007] and [8,0.18,0.009] respectively. The training was carried out in a Jupyter notebook using an NVIDIA GeForce RTX 2060 SUPER GPU with 8GB of memory and an AMD Ryzen 7 3700X 8-Core CPU.

The first segmentation network was trained on three full MBM image taken from three different sources functionalized with laminin for sickle red blood cell adhesion. The chosen images originally had dimensions of 15171×2391, 15171×4782, and 5057×1196 pixels respectively. When creating the masks for this segmentation network, pixels corresponding to the adhered category were manually colored white. After the manual labeling was completed, the masks were sent through a python code which labeled all background pixels as black. The number of training/validation tiles generated for this segmentation network was 2163 tiles. Augmentations were applied to the training tiles, resulting in 17304 total tiles for training/validation/testing. 70 percent of the tiles were used for training, 20 percent were used for validation, and 10 percent were used for testing. When splitting the tiles into training/validation/testing groups, the training set was drawn from only one of the three image sources, while the validation/testing sets were drawn from the other two image sources. Furthermore, the training/validation/tesing datasets were stratified, to make sure that the relative distributions of classes were equal for all datasets.

The second segmentation network was trained on tiles from 30 frames of MBM videos extracted from three separate sources of CAR-T cell adhesion to E-selectin. Ten frames from each source were used, where the ten frames were chosen from the last 100 frames of the video in increments of ten. Each frame had dimensions of 1392×1040 pixels. When creating the masks for this segmentation network, pixels corresponding to the adhered category were manually colored green. After the manual labeling was completed, the masks were sent through a python code which labeled all background pixels as black. These frames generated a total of 767 training/validation /testing tiles, where 70 percent were used for training, 20 percent for validation, and 10 percent for testing. When splitting the tiles into training/validation/testing groups, the training set was drawn from only one one video source, while the validation/testing sets were drawn from the remaining two sources. Augmentations were applied to the training/validation/testing tiles, resulting in a total of 6136.

### MBM with automated analysis with multiple cell types

When multiple cell types are expected to be visible in an MBM image, phase 2 classification becomes more complicated. In general for this case using a simple physical parameter (like cell size / eccentricity) for groups of adhered pixels will not be sufficient for accurate classification. The reason for this is because the distribution of physical parameters can overlap between the different types. Thus, trying to classify with only a physical parameter may lead to errors. One solution to this problem is using a machine learning classification network for phase 2, which we demonstrate here as a proof of concept to classify CAR-T cell adhesion to P-selectin in the presence of red blood cells (where both CAR-T and red blood cells are expected to adhere).

## Results

For simplicity, we re-used the phase 1 segmentation network from analysis of sickle red blood cell adhesion to laminin for analysis of CAR-T cell adhesion to Pselectin. To demonstrate the necessity of a machine learning approach for phase 2, we show simple physical characteristics (eccentricity versus size) of the adhered pixel groups in Fig. SI2. As in main text Fig. 4, there are only two distinct peaks: one at small areas (misclassified pixels / detritus) and one at large areas (all adhered cells). The two types of adhered cells do not form distinguishable peaks.

To get reliable CAR-T cell classification, we trained a ResNet50 convolutional neural network^44^ as described in the next section. The goal was to identify adhered pixel groups as either CAR-T cells or as non-CAR-T (“other”), starting with pixel groups that had sufficiently large areas to be cells (i.e. after passing the size threshold). The output of this binary classification network is a scalar between 0-1, which can be roughly thought of as the probability of the group being a CAR-T cell. To positively assign a CAR-T classification, we have a choice of confidence threshold, with all network outputs above the threshold labeled as CAR-T. The default threshold would be 0.5, but higher thresholds can also be used to get more conservative assignments. The effectiveness of the binary classification can be measured via an *F*_1_ score, which is harmonic mean of precision and recall (with values in the range 0-1, where 1 corresponds to perfect classification). As shown in Fig. SI3a, most confidence thresholds in the range 0.5-0.99 (except the highest ones) give excellent *F*_1_ scores *≳* 90% for classifying a set of 115/770 CAR-T/other cells. We chose a threshold of 0.67, and for this value Fig. SI3b shows the corresponding confusion matrix and Fig. SI3c shows a comparison to manual counting of CAR-T cells by human experts. The excellent agreement (*R*^2^ = 0.95) shows that is possible to have accurate phase 2 classification of MBM images via neural networks even in the presence of multiple cell types.

### Phase 2 training details

The classification network is a ResNet50 architecture pretrained on ImageNet^45^ with additional trainable layers added at the end (Fig. SI4). Note there is a special pre-processing applied to MBM images for CAR-T cell adhesion to P-selectin. Each pixel from each MBM image is color adjusted according to the equation:

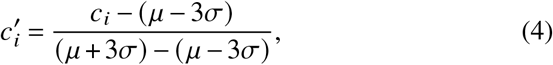

where 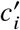 is the color of a pixel after adjustment, *c*_*i*_ is the original color of the pixel, *μ* is the mean color of the MBM image, and *σ* is the standard deviation of the color of the MBM image. This color adjustment is not strictly necessary, but it helps the phase 1 segmentation network generalize to a new experimental set-up (the alternative would have been to re-train the phase 1 network for this experimental case).

To train the classification network, 40*x*40 pixel regions are extracted from MBM images. These regions are centered on groups of adhered pixels classified by the phase 1 segmentation network which have areas larger than the size threshold of 180 pixels. To create a training/validation dataset these regions are manually classified as CAR-T or other. We extracted 8830 regions, and classified 1143 CAR-T cells, and 7687 other, where 70% of regions were used for training, 20% of images were used for validation, and 10% of regions were used for generating *F*_1_ scores and confusion matrices. Data augmentations such as horizontal and vertical flips were applied to the training/validation data set, resulting in four unique orientations for each image, and giving 3200/21523 CAR-T/other images for training and 914/6149 CAR-T/other images for validation. Class weighting was used for all phase 2 training according to the ratios of the classes relative to the total number of images. A stop function was included in the training, which automatically stopped the training after the validation loss had not decreased for 6 straight epochs. The best weights of the network after the 6 straight epochs were used as the trained network weights. The training was carried out via an Adam optimizer with a learning rate of 0.00035 and batch size of 32. The loss function used for optimization was a categorical cross-entropy function from the Keras library. The training was completed on a Jupyter notebook using an NVIDIA GeForce RTX 2060 SUPER GPU with 8GB of memory and an AMD Ryzen 7 3700X 8-Core CPU.

### Hyperparameter Optimization of Machine Learning Training

Using the Hyperopt Python package (based on the Tree-Parzen search algorithm), we were able to optimize the hyperparameters for both phase 1 and phase 2 of the ML pipeline. When training phase 1 segmentation networks, we focused on three hyperparameters: the learning rate, the patience (how many epochs we wait without improvement to stop training), and the *α* parameter from our loss function. For phase two training, we also had three hyperparameters: the learning rate, the patience, and the model architecture. To optimize the hyperparameters in phase 1 and phase 2, the networks were repeatedly trained, while the Hyperopt package suggested parameters in the space. The spaces defined were; [0.0001-0.01] for the learning rate, [0-10] for the patience, [0,1] for the *α* parameter of phase 1 training, and [VGG16 vs. ResNet50 vs. EfficientNetB3] for the model architecture for phase 2 training. The networks were optimized on the objective of minimizing the Jaccard-Loss on the testing datasets. Once the Jaccard-Loss did not decrease for ten straight parameter samplings, and after at least 50 samplings were made, the optimization was stopped.

**Fig. SI1.**
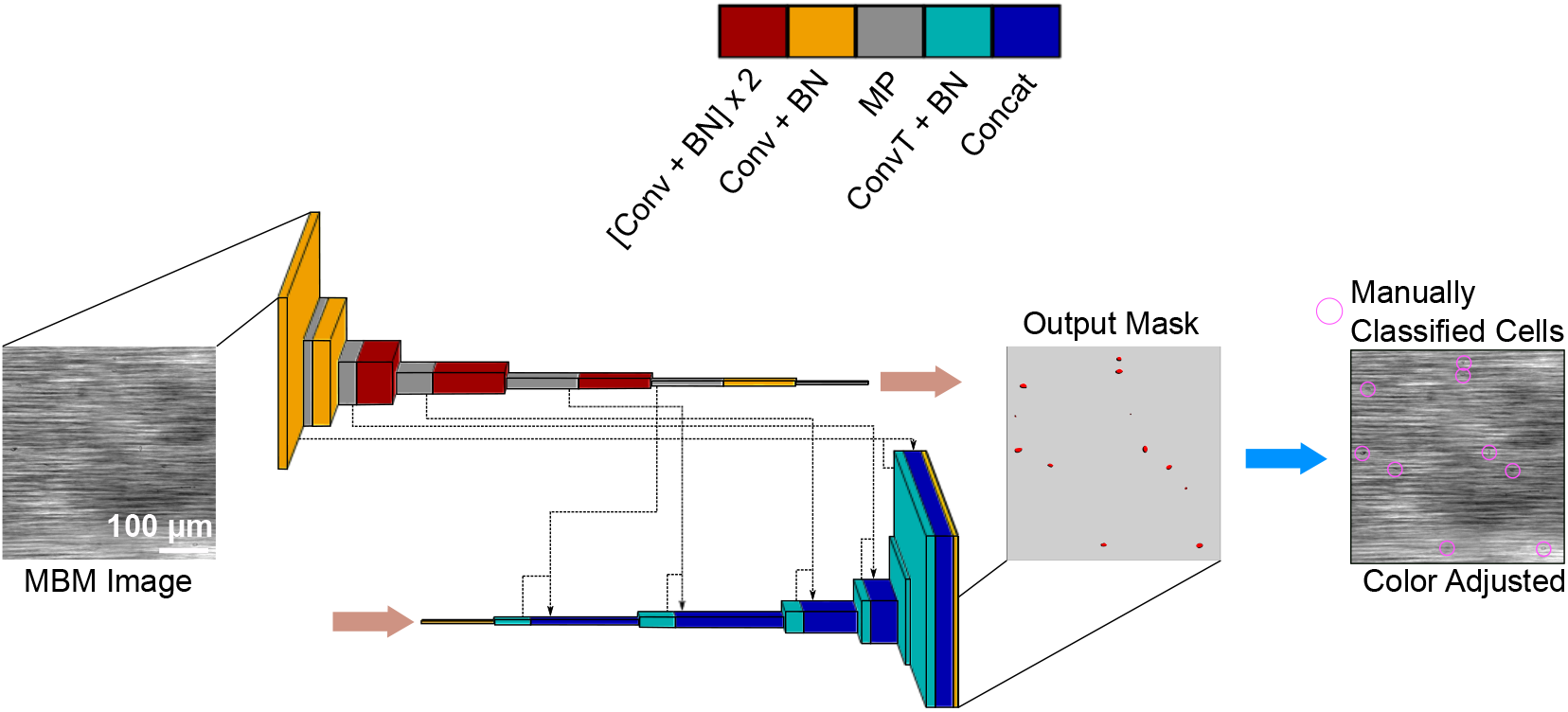
The architecture of the phase 1 segmentation network. The architecture is a modified U-Net, based on Ref. 43. We show an example input and output for sickle red blood cell adhesion to laminin, as well as manual classification for comparison. The groups of adhered pixels found by the segmentation network appear to match manual classification well. Conv = 2D Convolutional. BN = Batch Normalization. MP = Max Pooling. ConvT = 2D Convolutional Transpose. Concat = Concatenation.

**Fig. SI2.**
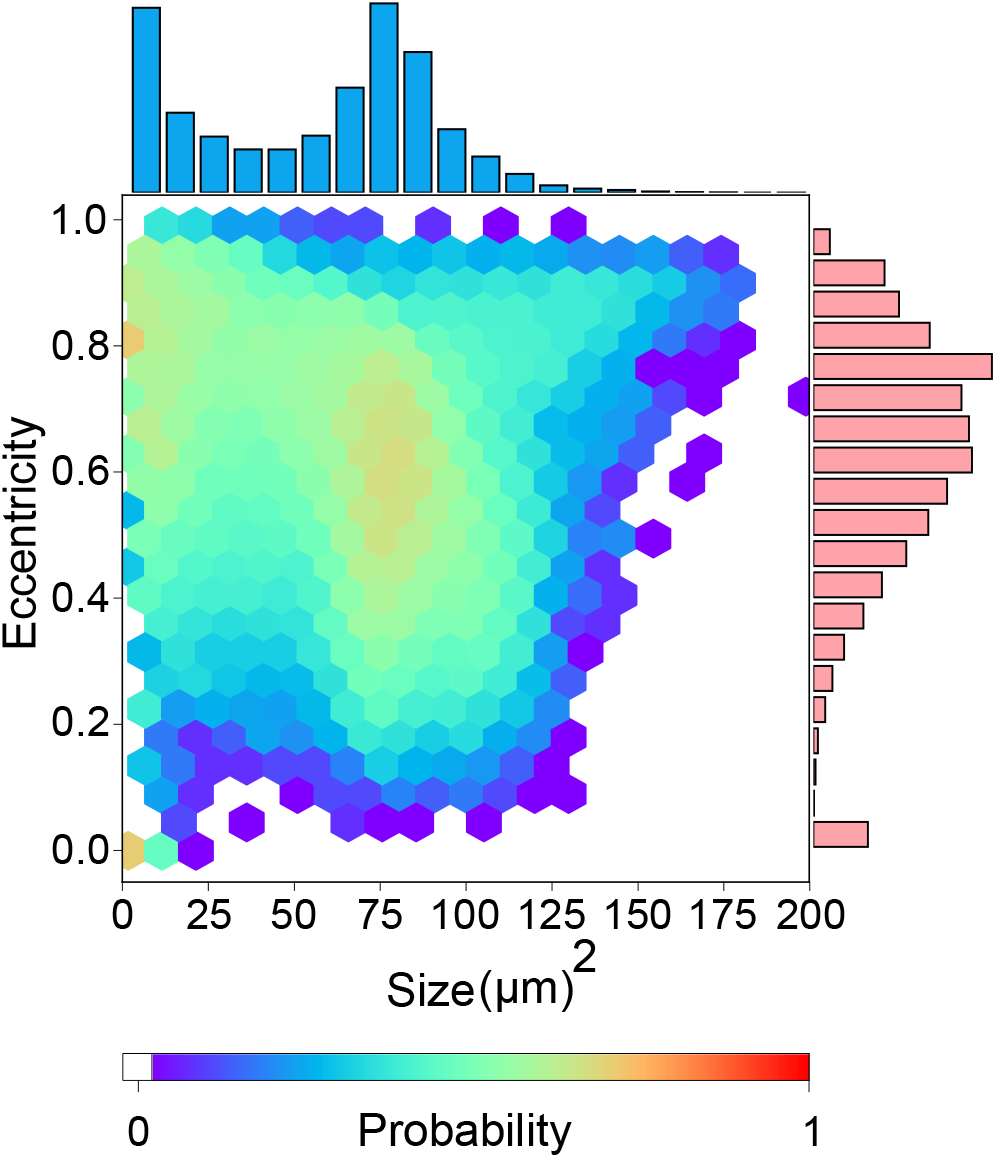
CAR-T cell/sickle red blood cell adhesion to P-selectin size vs. eccentricity distribution. Hexplot showing the joint distribution of size and eccentricity for CAR-T/red blood cell adhesion to P-selectin for 947 consecutive video frames (*N* = 82572). Projections on each axis show the marginal distributions of size and eccentricity alone. Note that the distribution peak at large areas corresponds to both CAR-T and red blood cells, and so the two types cannot be distinguished via size/eccentricity characteristics alone.

**Fig. SI3.**
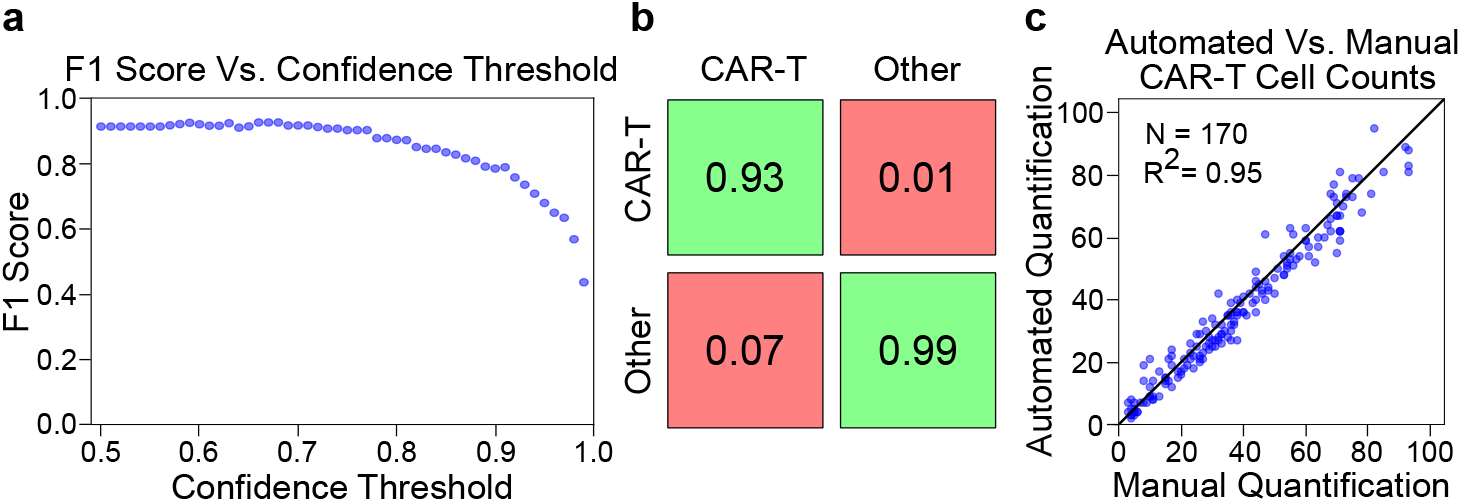
Validation of MBM with automated analysis with more than one cell type. **a)** *F*_1_ score vs. confidence threshold plot. There are a range of confidence thresholds which produce acceptable *F*_1_ scores. **b)** Confusion matrix for the classification network on a subset of groups of adhered pixels (*N* = 885), for a confidence threshold of 0.67. The true positive rate is large for quantified cells. **c)** Comparison between automated and human counts of CAR-T cell adhesion to P-selectin. The *R*^2^ value indicates good agreement between the two counts.

**Fig. SI4.**
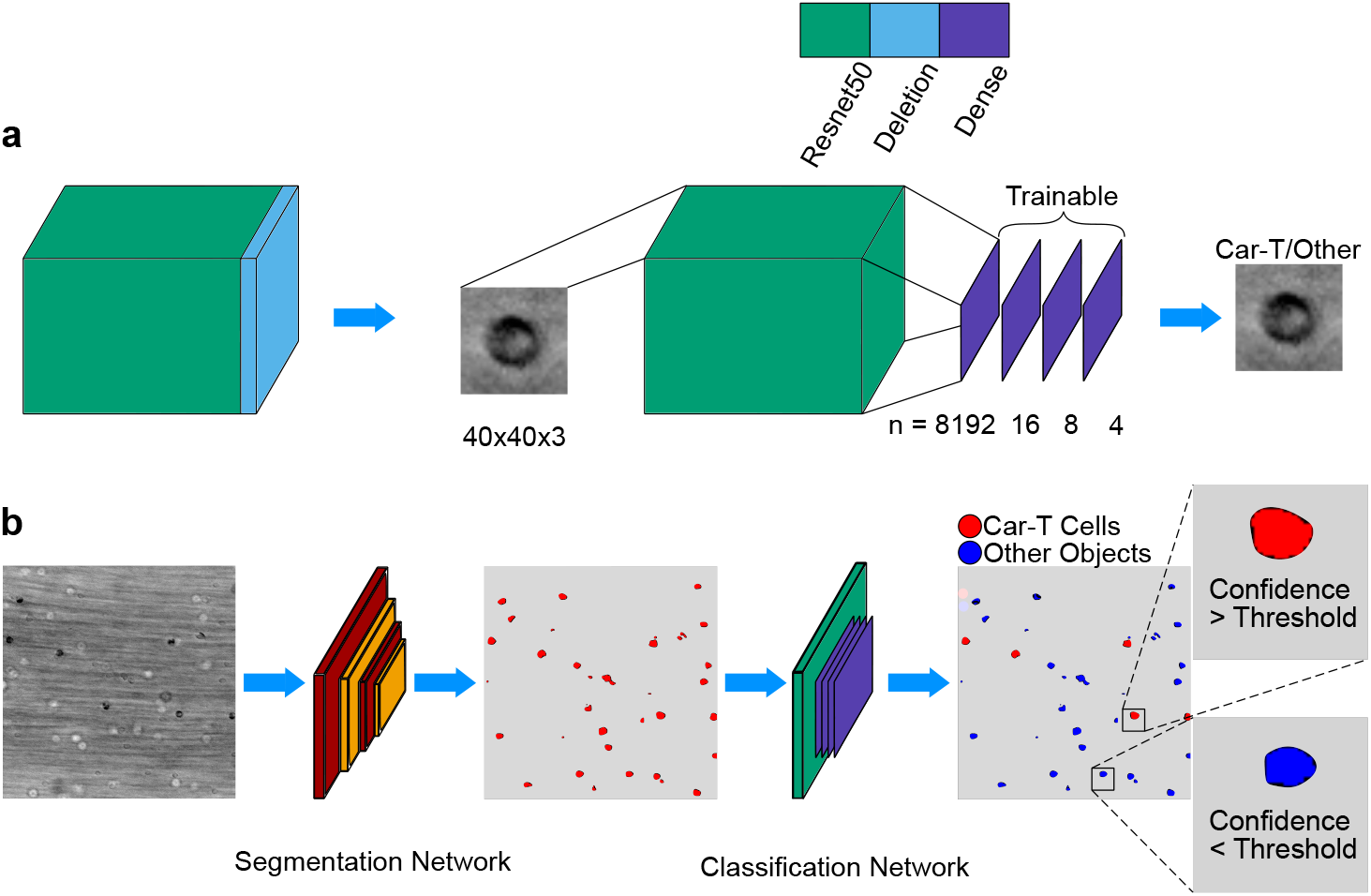
Phase 2 classification network architecture. **a)** Schematic of the architecture for the phase 2 classification network. Only the final four layers are trainable. Conv = 2D Convolutional. MP = Max Pooling. D = Dense. **b)** Cartoon representation of MBM with automated analysis for analysis of multiple cell types. Classification now requires a neural network.

**Fig. SI5.**
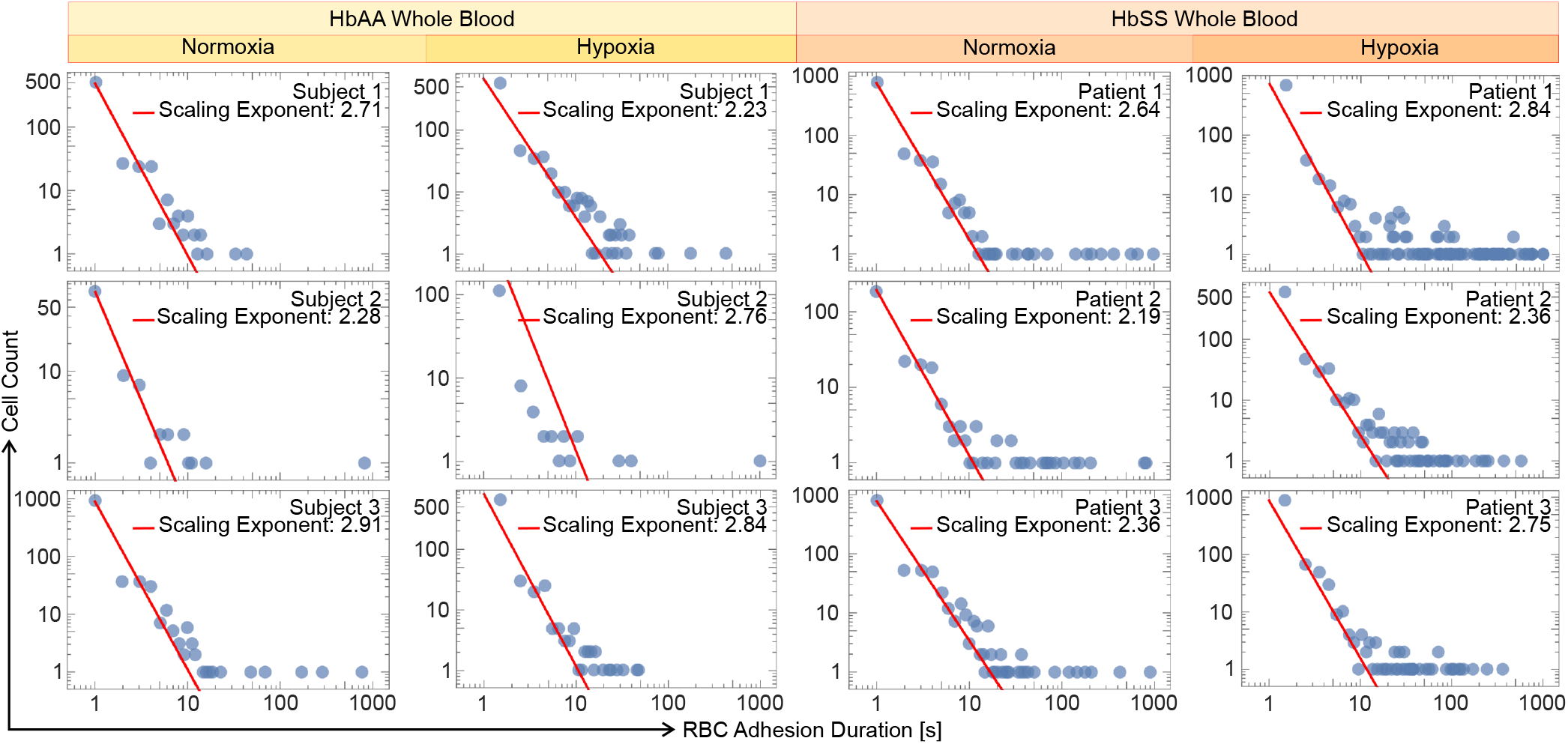
Adhesion duration profiles for red blood cells (RBCs) with HbAA and HbSS genotype, and under normoxic and hypoxic conditions during whole blood flow. The adhesion duration of RBCs shows a power law distribution at short times (red lines indicate estimated scaling exponents). HbSS-containing sickle RBCs establish longer-lasting bonds more than HbAA-containing RBCs, but short-duration adhesion events show a resemblance between genotypes. Hypoxia increases the long-duration adhesion events in only sickle whole blood samples.

**Fig. SI6.**
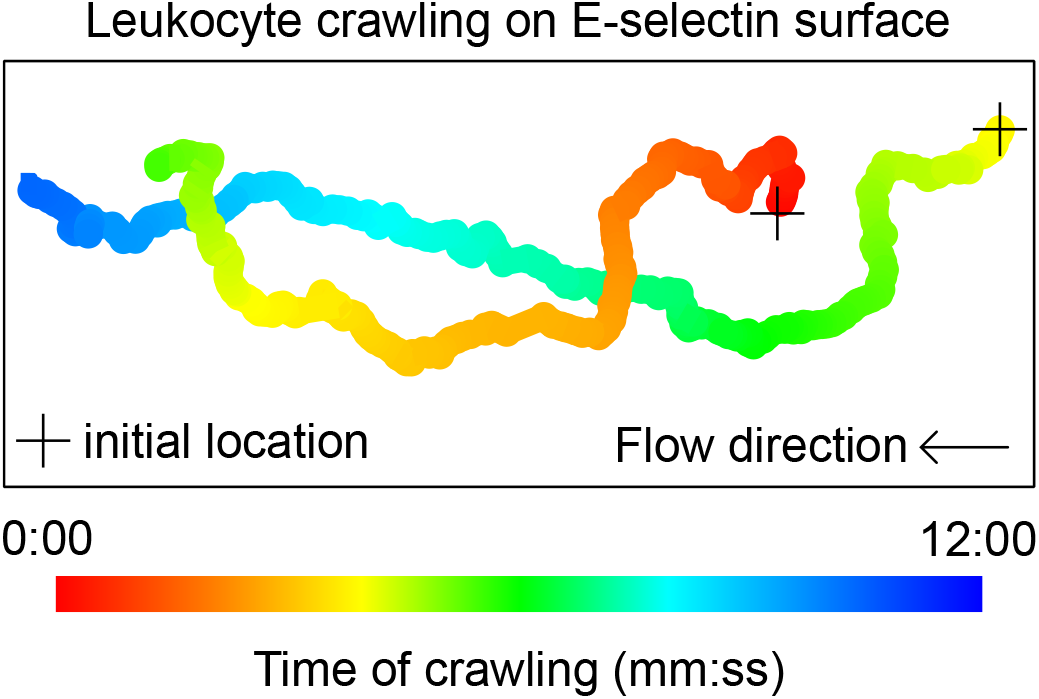
Paths of two crawling leukocytes on an E-selectin surface under whole blood flow. We used whole blood from a healthy subject collected in a heparin tube and we did not perform any preprocessing. Path color denotes how much time is spent during crawling. The initial locations shown are random.

**Fig. SI7.**
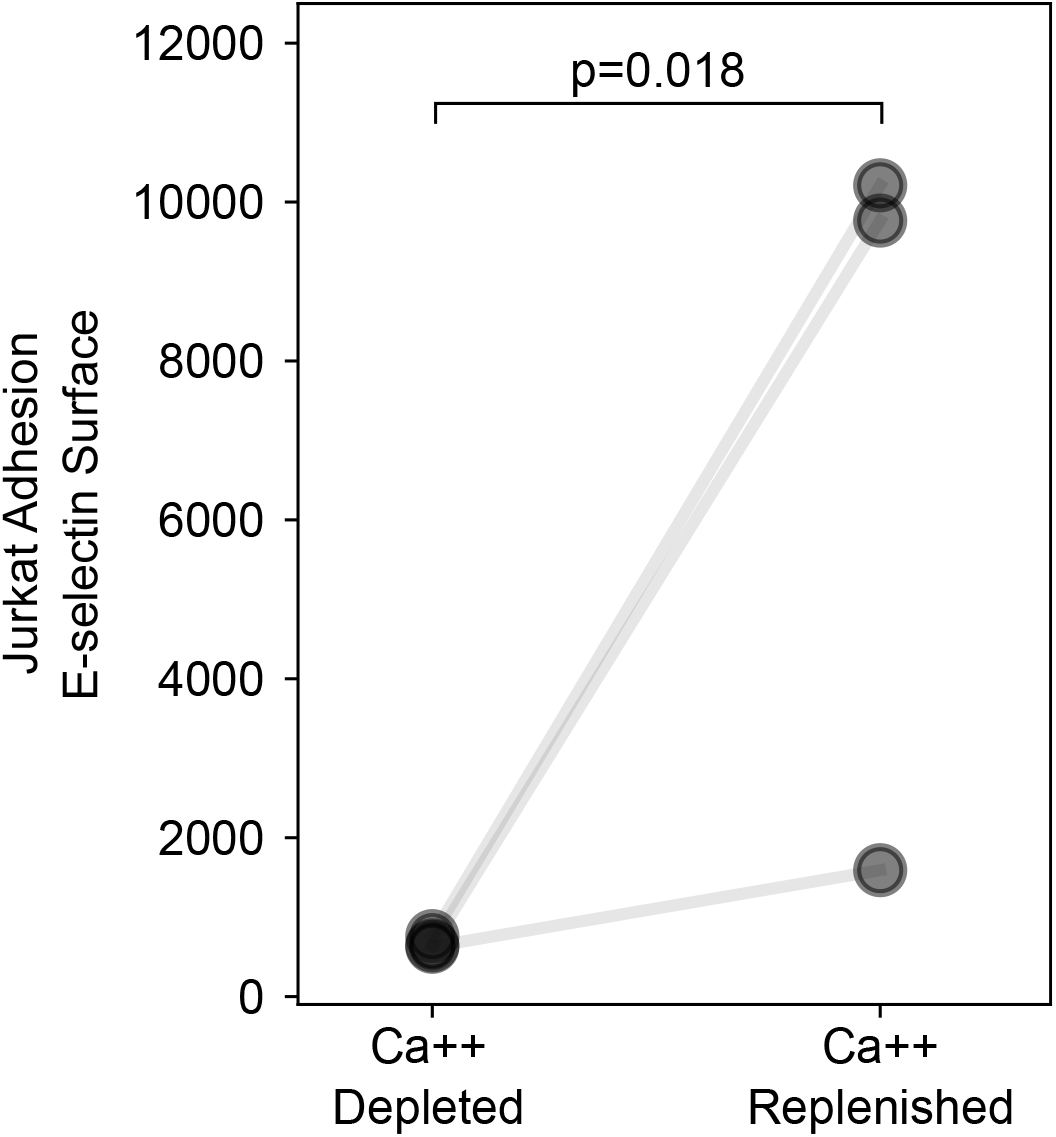
Replenishing Ca++ increases the number of Jurkat cells adhered to E-selectin significantly. Blood sample collection with EDTA tubes prevents coagulation by calcium chelation, but calcium is crucial for leukocyte activity. Ca++ is replenished by resuspending the mixture of HbAA RBCs and Jurkats at 40% hematocrit in Hank’s buffer containing calcium. Control is resuspension in PBS without calcium.

**Video SI1.**
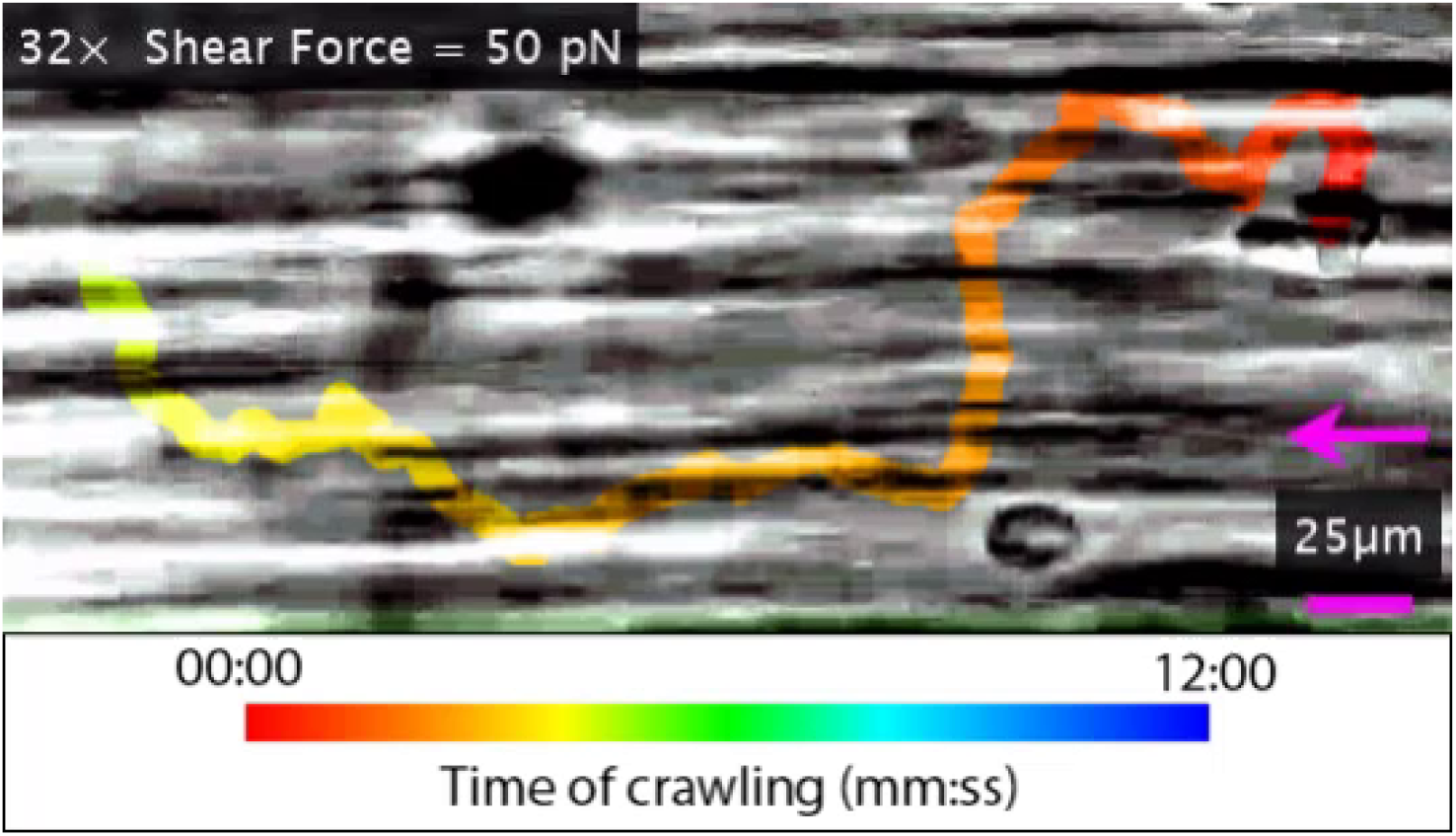
Video of a crawling leukocyte on E-selectin surface under whole blood flow. The blood sample was collected in a heparin tube from a healthy subject and tested immediately after collection. Video is fast forwarded 32 times, and color indicates the time of the crawling.

